# Photoactivation of silicon rhodamines via a light-induced protonation

**DOI:** 10.1101/626853

**Authors:** Michelle S. Frei, Philipp Hoess, Marko Lampe, Bianca Nijmeijer, Moritz Kueblbeck, Jan Ellenberg, Jonas Ries, Stefan Pitsch, Luc Reymond, Kai Johnsson

## Abstract

We present a new type of photoactivatable fluorophore that forms a bright silicon rhodamine derivative through a light-dependent isomerization followed by protonation. In contrast to other photoactivatable fluorophores, no caging groups are required, nor are there any undesired side-products released. Using this photoactivatable fluorophore, we created probes for HaloTag and actin for live-cell single-molecule localization microscopy and single-particle tracking experiments. The unusual mechanism of photoactivation and the fluorophore’s outstanding spectroscopic properties make it a powerful tool for live-cell super-resolution microscopy.

## Introduction

Advances in super-resolution microscopy (SRM) have led to unprecedented insights into cellular structures and processes over the past decade^1–2^. One of these SRM approaches is single-molecule localization microscopy (SMLM), which relies on the switching of fluorophores between an ‘off’ and an ‘on’ state^3–5^. The switching can be achieved by using photoactivatable or switchable fluorophores^6–8^. As small-molecule fluorophores are generally brighter and more photostable than fluorescent proteins^9^, they are of advantage for SMLM experiments^10–11^. Photoactivatable (or caged) small-molecule fluorophores are known throughout many of the different fluorophore families and are mainly synthesized using photolabile protecting groups^11–14^. Photoactivatable rhodamine derivatives have been obtained through the attachment of *ortho*-nitrobenzyl moieties^11^. However, these probes are mostly used in fixed-cell microscopy due to their decreased solubility and poor cell-permeability^15–17^. Furthermore, they result in the stoichiometric formation of very electrophilic nitroso-aldehydes or ketones as reactive byproducts, which are toxic and of concern in live-cell imaging^18^. Rhodamines have also been rendered photoactivatable through a diazoketone group^19^, leading to the introduction of the photoactivatable Janelia Fluor dyes PA-JF_549_ and PA-JF_646_^20^, which have been successfully used for fixed-cell and live-cell SMLM. However, photoactivation of these fluorophores leads to the formation of a dark side-product. The extent, to which the undesired side-product is formed, depends on the structure and environment of the fluorophore complicating applications of the diazoketone approach. In addition, photoactivation of fluorophores caged with the diazoketone group proceeds through a carbene, which can react with intracellular nucleophiles^21^. In light of the limitations of the existing caging strategies, alternative chemical strategies are needed to generate photoactivatable fluorophores. Here, we report the discovery, synthesis and characterization of a new class of cell-permeable, photoactivatable fluorophores (PA-SiRs), which are based on the silicon rhodamine (SiR) scaffold and activated through an unprecedented light-induced protonation. We demonstrate the utility of these fluorophores for live-cell SMLM of intracellular targets and single particle tracking experiments.

## Results and Discussion

### Synthesis and *in vitro* Characterization

The first analogue of this new class of fluorophores was serendipitously found during the attempted synthesis of a SiR derivative bearing an alkyl chain in place of the aromatic substituent at the 9 position of the xanthene scaffold (Fig. 1a). Instead of the desired fluorescent SiR **2** we isolated the non-fluorescent isomer **PA-SiR** (**1**). **PA-SiR** possesses an exocyclic double bond and the two aromatic ring systems are not conjugated, reflected by its λ_abs, max_ value of 290 nm. However, **PA-SiR** underwent isomerization and protonation upon UV irradiation in aqueous solution, re-establishing the fluorescent xanthene core of SiR **2** (Fig. 1a, b and Supplementary Fig. S1, 2). The photoproduct SiR **2** showed an absorption maximum at λ_abs, max_ = 646 nm and emitted at around 660–670 nm. Its extinction coefficient of ε_646_ = 90’000 ± 18’000 m^−1^ cm^−1^ and fluorescence quantum yield φ = 19.0 ± 2.4% in aqueous buffer were only marginally smaller than those of the previously described SiR-carboyxl^22^ (Fig. 1b and Supplementary Table S1). However, **2** is susceptible to nucleophilic attack by water leading to rapid establishment of an equilibrium between **2** and **3** (Fig. 1a, c, e and Supplementary Fig. S3). Structural modifications on **PA-SiR** can influence this equilibrium as demonstrated by several synthetized analogues (Supplementary Fig. S4). Moreover, both photoactivation of **PA-SiR** as well as the equilibrium between **2** and **3** are pH sensitive (Fig. 1d and Supplementary Fig. S5). Photoactivation is prevented by protonation of the aniline groups and is therefore highest at pH values above pH = 6 as revealed by measuring the maximal absorbance at 646 nm reached directly after activation (A_max_). The equilibrium between **2** and **3**, as measured by recording the absorbance at equilibrium and correcting for A_max_ at 646 nm (A_eq_), was shifted towards **3** at higher pH values (Fig. 1d). At physiological pH only about 10% of the activated **PA-SiR** was present as SiR **2** in comparison to 80% at pH = 6.1. Noteworthy is also the quantitative nature of the photoconversion of **PA-SiR**, which becomes apparent when following the conversion of **PA-SiR** to **3** by nuclear magnetic resonance (NMR). These experiments also revealed that the photoactivation is reversible on a time scale of days (Fig. 1e and Supplementary Fig. S3).

**Figure 1:**
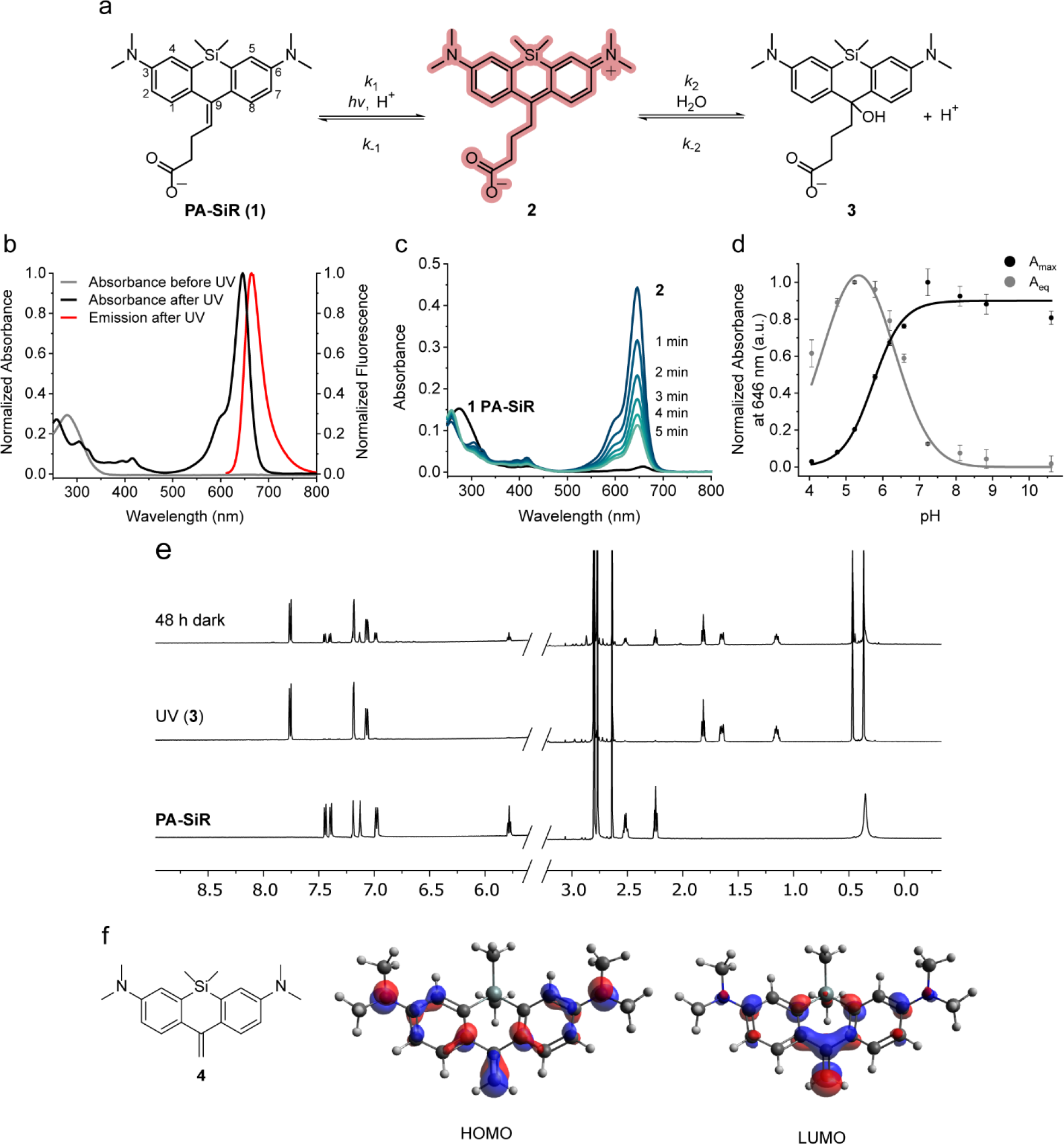
Structure and properties of PA-SiR. **a**, Reaction scheme for photoactivation of **PA-SiR** (**1**), and equilibrium between **2** and **3**. **b**, Normalized absorption spectra of **PA-SiR** in PBS (10 μM) before and after UV irradiation as well as emission spectra after activation. **c**, Absorption spectra of **PA-SiR** in PBS (10 μM) before activation and directly after UV irradiation measured every 1 min, revealing the reaction from **2** to **3**. **d**, pH dependence of the equilibrium system of **PA-SiR** in PBS (10 μM) at different pH after brief photoactivation through UV irradiation. Normalized absorbance values A_max_ directly after activation and A_eq_ in equilibrium at different pH values are given, reflecting changes in activation (A_max_) and equilibrium constant (A_eq_). Values displayed are means from three individual measurements, error bars correspond to 95% confidence intervals. **e**, ^1^H nuclear magnetic resonance (NMR) spectra of **PA-SiR** (2.0 mM in PBS) before UV irradiation, after complete conversion to **3** and after further 48 h in the dark. (For assignment of peaks see Supplementary Fig. S3). **f**, Structure of model PA-SiR **4** together with calculated HOMO and LUMO (B3LYP/6-31G(d), only contributions bigger than 0.05 are shown). According to these calculations, the LUMO receives dominant contributions from the exocyclic double bond whereas the HOMO locates to the anilines. This indicates that a light-induced HOMO-LUMO transition would lead to an intramolecular charge transfer.

To the best of our knowledge, this type of light-induced protonation has not previously been reported for rhodamine derivatives or other xanthenes (see Supplementary Fig. S1 for an overview of related reactions). Calculations of the frontier molecular orbitals of model compound PA-SiR **4** and data published on cross-conjugated 1,1-diphenyl alkenes^23–24^ suggest that the photoactivation proceeds through a twisted intramolecular charge transfer followed by protonation of the intermediate (Fig. 1f, Supplementary Fig. S1).

The susceptibility of activated **PA-SiR** towards nucleophiles and its half-life of minutes at physiological pH are a disadvantage of these fluorophores for standard diffraction limited imaging. However, this is less relevant for single-molecule based super-resolution microscopy since the observation period of individual fluorophores in SMLM is on the order of milliseconds and the reaction of activated **PA-SiR** with nucleophiles should not interfere in such experiments. Furthermore, the equilibrium of the reaction of activated **PA-SiR** with nucleophiles is environmentally sensitive. In fact, when we prepared conjugates of **PA-SiR** with ligands for protein labeling (Supplementary Fig. S6, S7)^25–28^, we discovered that **PA-SiR-Halo** attached to HaloTag can be very efficiently activated and its fluorescent form **2** is stable over hours at physiological pH, whereas **PA-SiR-Halo** not conjugated to HaloTag is inefficiently activated and the activated probe decays quickly (Fig. 2a-b, Supplementary Table S1 and Supplementary Fig. S7). This apparent fluorogenicity of the probe should prove beneficial for live-cell imaging as unconjugated **PA-SiR-Halo** is not fluorescent, which increases the signal-to-background ratio. Additionally, **PA-SiR-Halo** conjugated to HaloTag and photoactivated showed much greater stability towards other nucleophiles such as cysteamine than free **PA-SiR** (Fig. 2c). The generated fluorescent product had an extinction coefficient of ε_646_ = 180’000 ± 30’000 M^−1^ cm^−1^ and a fluorescence quantum yield of φ = 29.2 ± 1.2% in aqueous buffer making it an outstanding fluorophore. Its quantum yield of activation was found to be φ_act_ = 0.86 ± 0.07% at 340 nm, similar to that of photoactivatable proteins (Supplementary Table S1)^29^.

### *In cellulo* Characterization

**PA-SiR-Halo** possesses a number of properties that make it an attractive candidate for live-cell imaging such as (i) the absence of side-products during photoconversion, (ii) the absence of caging groups that affect solubility and permeability, (iii) the efficiency of photoactivation and stability of the HaloTag-bound probe compared to unconjugated probe, and (iv) its outstanding spectroscopic properties. We therefore incubated U-2 OS cells expressing a histone H2B-HaloTag fusion protein with 0.5 μM **PA-SiR-Halo** for 2 h and imaged the cells prior and after UV activation at 365 nm (Fig. 2d, e). We found that **PA-SiR-Halo** showed an excellent signal-to-background ratio after activation under no wash conditions (32 ± 5, mean ± 95% confidence interval, *N* = 119 cells) and that the fluorescence signal after activation was stable over time. In comparison, **PA-JF_646_-Halo** showed faster activation kinetics but a lower signal-to-background ratio after activation (13.2 ± 1.9, *N* = 121 cells) (Supplementary Fig. S9d-f)^20^. Moreover, **PA-SiR-Halo** was used to image various other intracellular HaloTag fusion proteins (Fig. 2f-h). Taken together, these experiments validate that **PA-SiR-Halo** is suitable for live-cell imaging. It should be noted that other **PA-SiR** probes can be generated (Supplementary Fig. S6, S7). Specifically, we attached **PA-SiR** to the F-actin-binding natural product jasplakinolide, yielding **PA-SiR-actin**, and used it successfully for live-cell imaging of actin filaments (Supplementary Fig. S7b, S9c)^27–28^.

**Figure 2:**
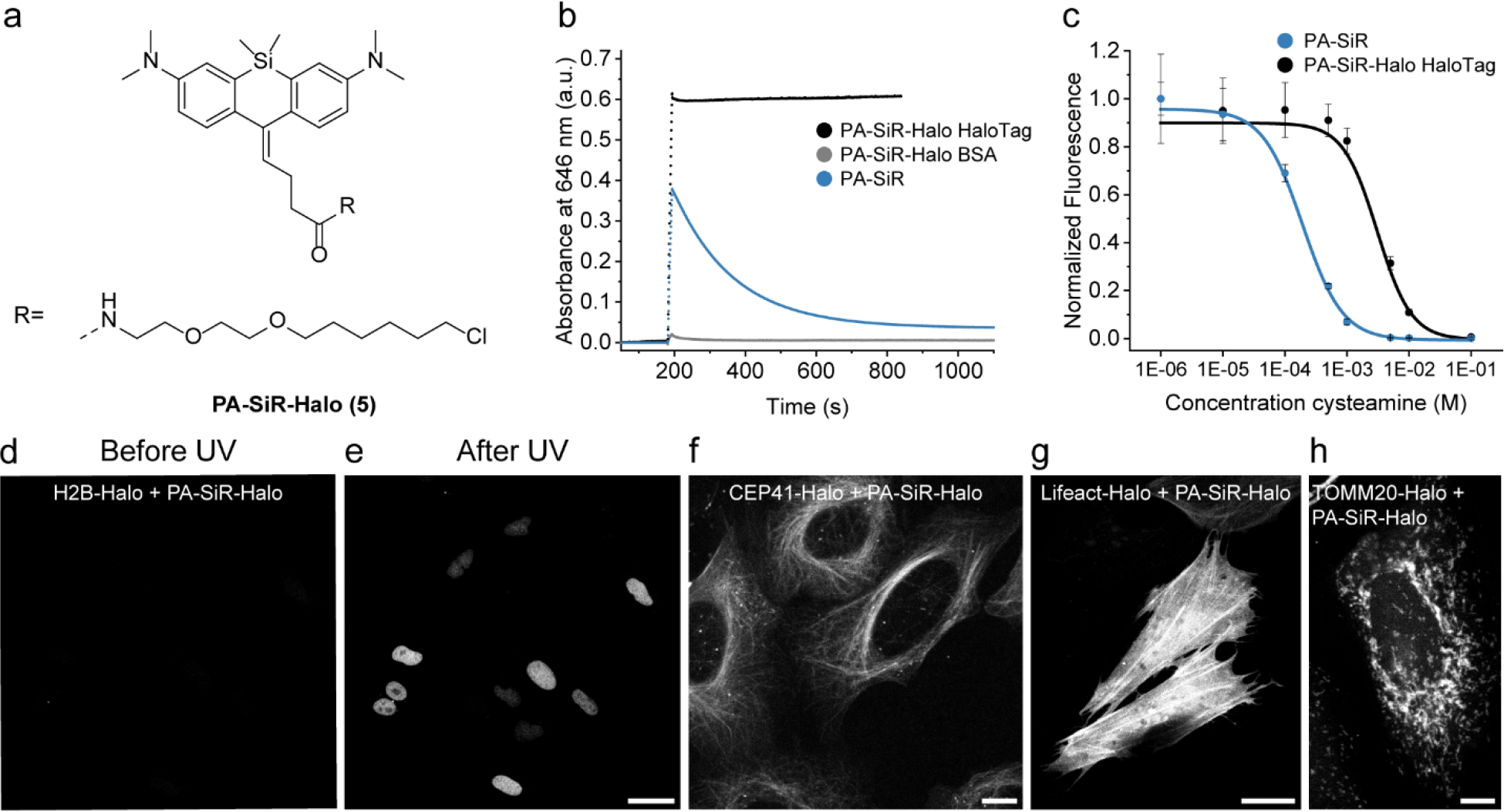
PA-SiR-Halo and the influence of HaloTag on its equilibrium system. **a**, Chemical structure of **PA-SiR-Halo** (**5**). **b**, Absorbance measurements at 646 nm over time for **PA-SiR** and **PA-SiR-Halo** in PBS (10 μM). **PA-SiR-Halo** was measured with addition of BSA or HaloTag (20 μM). **c**, Fluorescence signal after addition of cysteamine (0.001 – 100 mM) to fully activated **PA-SiR** or **PA-SiR-Halo** on HaloTag solutions in equilibrium (1 μM dye on 2 μM HaloTag). The effective concentrations at which half maximal fluorescence intensity was reached (EC_50_ values) were determined to be 0.192 ± 0.019 mM for **PA-SiR** and 3.1 ± 0.5 mM for **PA-SiR-Halo** (mean ± 95% confidence interval, both *N* = 24 samples). **d-e**, Maximum projection of a z-stack of U-2 OS cells stably expressing H2B-Halo stained with **PA-SiR-Halo** (0.5 μM for 2 h) before **d** and after UV irradiation **e**. Scale bar, 40 μm. **f-h**, Confocal images of several HaloTag fusion proteins stained with **PA-SiR-Halo** (0.5 μM for 1.5 h): **f**, microtubule binding protein CEP41-Halo. Scale bar, 10 μm. **g**, F-actin binding peptide LifeAct-Halo. Scale bar, 20 μm. **h**, TOMM20-Halo located in the outer membrane of mitochondria. Scale bar, 10 μm.

**Figure 3:**
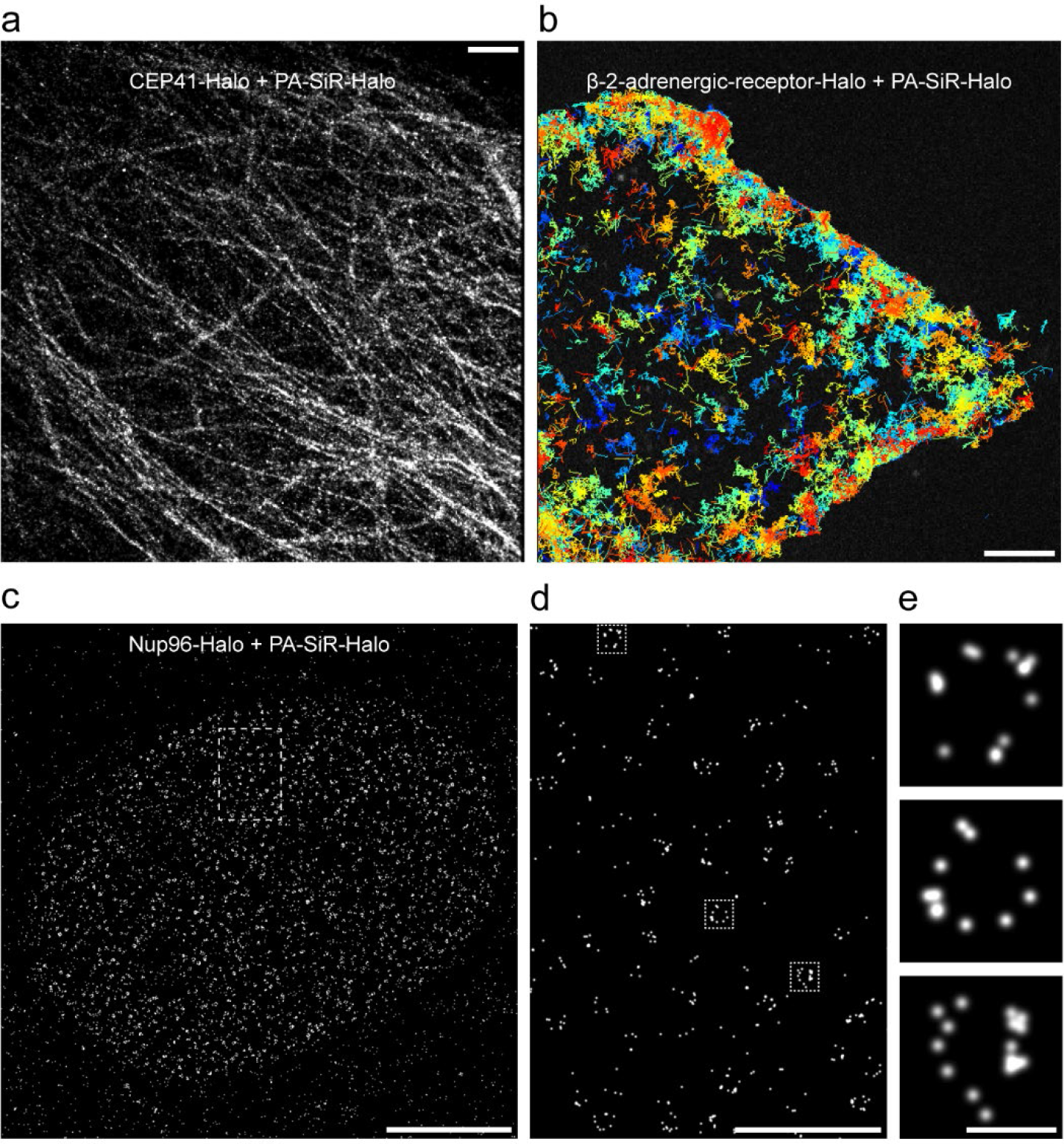
Super-resolution microscopy and single-particle tracking experiments. **a**, Super-resolved image of fixed U-2 OS cells stably expressing CEP41-Halo stained with **PA-SiR-Halo** (1 μM for 2 h). The image is reconstructed from 14’083 frames (100 ms exposure time, 2.9 kW cm^−2^ at 642 nm excitation). The average microtubule diameter was found to be 64 ± 10 nm (mean ± 95% confidence interval, *N* = 10 tubules). Scale bar, 1 μm. **b**, Image of cumulative single-particle tracks of β-2-adrenergic-receptor-Halo stained with **PA-SiR-Halo** (0.5 μM, 1 h) measured during 2 min. Continuous lines are drawn representing the movement of individual receptors. They are color coded in order to distinguish the individual tracks. For visibility, only tracks that have an overall displacement larger than 0.28 μm are shown (30 ms exposure time, 0.3 kW cm^−2^ at 642 nm excitation). Scale bar, 5 μm. **c**, Super-resolved overview image of endogenously tagged Nup96-Halo in U-2 OS cells stained with **PA-SiR-Halo** (1 μM for 2 h). Scale bar, 5 μm. **d**, Super-resolved image from the boxed region in **c**. Scale bar, 1 μm. **e**, Single nuclear pores from boxed regions in **d** following the same order (top-bottom). Scale bar, 100 nm.

### Fixed-Cell SMLM

The photophysical properties such as the number of detected photons per frame and fluorophore are decisive for SMLM as the attainable localization precision scales with the inverse square root of the number of detected photons^30^. In order to determine these numbers, we immobilized HaloTag labeled with **PA-SiR-Halo** on coated glass coverslips and imaged the fluorophore using total-internal reflection (TIRF) microscopy (Supplementary Fig. S10a). In these experiments, we used a 405 nm laser for photoactivation, generally used to create a sparse subset of fluorescent molecules in SMLM. We found that the photon numbers per particle per frame for **PA-SiR-Halo** at a power density of 1.2 kW cm^−2^ suitable for live-cell single-particle tracking were roughly 30% higher than for **PA-JF_646_-Halo** and considerably higher than those measured for mEOS3.2 (Supplementary Fig. S10b)^20^. To test the performance of **PA-SiR-Halo** in fixed-cell SMLM, we expressed the microtubule binding protein CEP41 as a HaloTag fusion in U-2 OS cells and labeled it with **PA-SiR-Halo**. The microtubule diameter was determined to be FWHM_**PA-SiR-Halo**_ = 64 ± 10 nm (mean ± 95% confidence interval, *N* = 10 tubules) which is consistent with previously found data (Fig. 3a, Supplementary Fig. S11a,d)^8^. Furthermore, we imaged a HaloTag fusion of NUP96^31^, a protein of the nuclear pore complex. Nuclear pores possess a regular circular shape with an internal diameter of about 100 nm^32–33^. Using **PA-SiR-Halo** labeled NUP96-Halo in fixed U-2 OS cells we were able to reveal the circular structure of the nuclear pore (Fig. 3c-e). In addition, **PA-SiR-Actin** was tested for SMLM in fixed COS-7 cells revealing stress fibers and connecting thinner fibers (Supplementary Fig. 11c). Both **PA-SiR-Halo** and **PA-SiR-Actin** are cell-permeable and make it possible to label live-cells, circumventing permeabilization steps during fixation and therefore reducing potential sources of artifacts^34^.

### Live-Cell Single-Particle Tracking and SMLM

We next tested the performance of **PA-SiR-Halo** in live-cell single-particle tracking photoactivated localization microscopy (sptPALM) (Fig. 3b)^35^. To this end, we chose to track a G-protein coupled receptor involved in cellular signaling that is located in the plasma membrane: beta-2-adrenergic receptor (β2AR)^36^. Using β2AR fused to HaloTag and labeled with **PA-SiR-Halo**, we were able to track β2AR for several hundreds of milliseconds before photobleaching (Fig. 3b). These track-lengths are considerably longer than what is commonly found for photoactivatable or photoconvertible proteins^20^ and similar to what we found for **PA-JF_646_-Halo**. Furthermore, β2AR labeled with either **PA-SiR-Halo** or **PA-JF_646_-Halo** moved with comparable mean speeds (Supplementary Figure S12). **PA-SiR-Halo** might prove to be beneficial over **PA-JF_646_-Halo** in intracellular single-particle tracking experiments, where high signal-to-background ratios are required.

Finally, we investigated the potential of **PA-SiR-Halo** for live-cell SMLM. In such experiments, we could follow the fast dynamics of mitochondria (TOMM20-Halo) labeled with **PA-SiR-Halo** over one minute in 10 s snapshots without artificial narrowing and collapsing of structures (Fig 4. and Supplementary Movie S1, Supplementary Fig. S13). The movie and the snapshots taken thereof revealed intermediate formation of thin tubules between mitochondria (blue arrowheads), as was previously seen with SMLM imaging of MitoTracker Red^37^. It was possible to follow fission events of mitochondria highlighting the dynamic network of connecting and disconnecting mitochondrial units (yellow arrowheads). Most interestingly, localizing the fluorophore to the outer membrane of the mitochondria further enabled us to distinguish the outer membrane from the matrix in several cases (red arrowheads), which has not been observed with live-cell SMLM so far. This will eventually help to study interactions between the inner and outer membrane of mitochondria by two color SMLM. This demonstrates that **PA-SiR-Halo** enables live-cell SMLM of intracellular targets.

**Figure 4:**
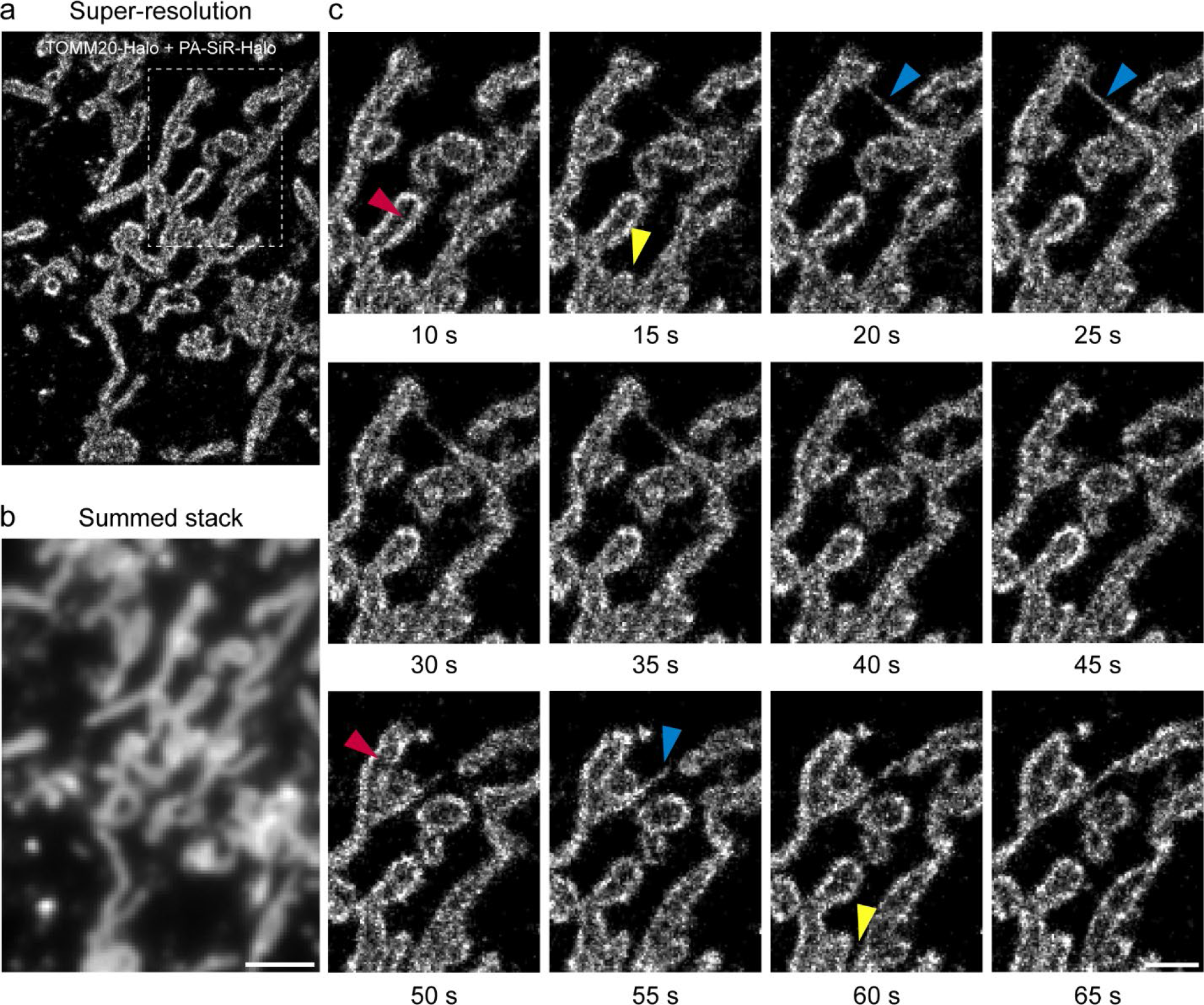
Live-cell super-resolution microscopy of TOMM20-Halo labeled with PA-SiR-Halo. **a**, Super-resolved image acquired within 10 s (50 ms exposure time, 0.3 kW cm^−2^ 642 nm excitation). The recently published ImageJ plugin HAWK^38^ was used to achieve imaging at high emitter densities to capture fast structural changes. **b**, Sum projection over the first 10 s mimicking the diffraction limited image. Scale bar, 2 μm. **c**, Time series of boxed region in **a**. Each frame is reconstructed from 200 frames (10 s). For clarity, snapshots are shown only every 5 s. Several mitochondria are perceived to be hollow as TOMM20 is localized to the outer membrane of mitochondria (red arrowheads). The highly dynamic mitochondria form thin tubules between neighboring mitochondria (blue arrowheads) and disconnect (fission) in other areas (yellow arrowheads). Scale bar, 1 μm. Full rolling frame movie available as supplementary movie S1.

## Conclusion

In summary, **PA-SiR** is a new type of photoactivatable, cell-permeable, far-red fluorophore that is activated by an unusual light-induced protonation. Its outstanding spectroscopic properties make it well suited for SMLM in both fixed and live-cells and enabled us to create powerful probes for HaloTag and actin. We expect that the exceptional properties of **PA-SiR** will be exploited in the future to create various other photoactivatable probes for live cell imaging.

## Acknowledgements

This work was supported by the Max Planck Society, the École Polytechnique Fédérale de Lausanne, a grant from the Swiss Commission for Technology and Innovation (CTI), the NCCR Chemical Biology, and the European Molecular Biology Laboratory (to P.H., M.L., B.N., M.K., J.E. and J.R.), the EMBL International PhD Programme (to P.H.), the European Research Council (ERC CoG-724489, to P.H. and J.R.), and the National Institutes of Health Common Fund 4D Nucleome Program (Grant U01 EB021223 / U01 DA047728 to J.E. and J.R.). All requests for the Nup96-Halo cell line should be directed to Jan Ellenberg.

The authors thank H. Farrants, Dr. J. Hiblot for sharing reagents, Dr. B. Koch for help with the establishment of the stable CEP41-Halo cell line, Dr. C. Sieben (EPFL) for valuable discussions and sharing of the Matlab analysis script, Dr. Rolf Sprengel (MPI for Medical Research) for the donation of the COS-7 cells, the electronic workshop of the Max Planck Institute for Medical Research for technical assistance, the NMR service of EPFL for assistance with the NMR experiments, and the Advanced Light Microscopy Facility (ALMF) at the European Molecular Biology Laboratory (EMBL) and Leica Microsystems for support.

## Author contributions

M.S.F., P.H., M.L., J.R., S.P., L.R. and K.J. planned the experiments and co-wrote the paper. L.R. made the first observation of **PA-SiR** photoconversion and originated the project. M.S.F. performed the chemical synthesis and characterization as well as the widefield and confocal measurements. M.S.F. performed the SMLM on CEP41-Halo, F-actin and mitochondria with assistance from M.L․. B.N., M.K. and J.E. provided the U-2 OS Nup96-Halo cell line. M.S.F. and P.H. performed SMLM on NUP96-Halo.

## Additional information

Correspondence should be addressed to L.R. or K.J. Requests for materials should be addressed to K.J.

