## Supplementary Information for "Photoactivation of silicon rhodamines via a light-induced protonation"

### Supplementary figures, table and movies inventory

**Supplementary Scheme S1.** Synthetic route to PA-SiR derivatives according to literature procedures 4
**Supplementary Figure S1.** Photoactivation reaction mechanism 28-29
**Supplementary Figure S2**. LC-MS analysis of the photoactivation 30
**Supplementary Figure S3.** $^{1}$H nuclear magnetic resonance (NMR) photoactivation experiments 31
**Supplementary Figure S4**. Absorbance measurements at 646 nm over time for different PA-SiR analogues (10 μM in PBS). 32
**Supplementary Figure S5.** Influence of different pH on the equilibrium system. 33
**Supplementary Figure S6.** Structures and absorption, excitation and emission spectra of PA-SiR derivatives. 34-35
**Supplementary Figure S7.** Absorbance measurements at 646 nm over time for different PA-SiR probes in PBS (10 μM). 36
**Supplementary Figure S8**. Saturation experiment of PA-SiRs at different concentrations. 37
**Supplementary Figure S9.** Confocal and widefield microscopy images. 38-39
**Supplementary Figure S10**. Single-molecule assay. 40
**Supplementary Figure S11.** Fixed-cell SMLM and quantification. 41-42
**Supplementary Figure S12.** Quantification of live-cell tracking experiments in U-2 OS cells. 43
**Supplementary Figure S13.** Live-cell SMLM experiment. 44
**Supplementary Table S1.** Spectral properties of different PA-SiR analogues. 45
**Supplementary Table S2**. Fit parameters from Gaussian fit in Fig. 1d. 46
**Supplementary Table S3**. Fit parameters from sigmoidal fit in Fig. 1d. 46
**Supplementary Table S4.** Averaged fit parameters from mono-exponential fits for PA-SiR probes in Fig. 2b and Supplementary Fig. S7a, b. 47
**Supplementary Table S5.** Fit parameters from the dose response fit in Fig. 2c and Supplementary Fig. S7c. 47
**Supplementary Table S6.** Averaged fit parameters from mono-exponential fits in Supplementary Fig. S4 and S5. 48
**Supplementary Table S7.** Fit parameters from mono-exponential fits of the third section for PA-SiR, PA-SiR-C3 (C3) and PA-SiR-C3-Halo (C3Halo) form the saturation experiment in Supplementary Figure S8. 49
**Supplementary Table S8.** Averaged fit parameters from the exponential fits for the first sections of the saturation experiments. 50
**Supplementary Table S9** Settings for the different microscopy experiments. 51-52
**Supplementary Movie S1.** SMLM rolling frame movie of mitochondria dynamics. 53

### Supplementary Methods

*Chemical Synthesis*

Materials and general information

All chemical reagents and anhydrous solvents for synthesis were purchased from commercial suppliers (Acros, Apollo, Armar, Bachchem, Biomatrik, Fluka, Fluorochem, LC Laboratories, Merck, Reseachem, Roth, Sigma-Aldrich and TCI) and used without further purification. Halo-NHBoc **23** and BG-$NH_2$ **26** were synthesized according to literature procedures[1-2]. Jasplakinolide-NHBoc **24** was obtained from a custom synthesis by Spirochrome AG. Composition of mixed solvents is given by volume ratio (v/v). Reactions in the absence of air and moisture were performed in oven-dried glassware under Ar or $N_2$ atmosphere. Flash column chromatography was performed using a CombiFlash Rf system (Teledyne ISCO) using $SiO_2$ RediSep® Rf columns at 25 °C or a Biotage (Isolera™) flash system using SiliaSep™ columns. The used solvent compositions are reported individually in parentheses. Analytical thin layer chromatography was performed on glass plates coated with silica gel 60 F254 (Merck). Visualization was achieved using UV light (254 nm). Evaporation *in vacuo* was performed at 25–60 °C and 900–10 mbar. $^1H$, $^{13}C$, and $^{19}F$ NMR spectra were recorded on AV 400, Ascend™ 400 and AV 600 Bruker spectrometers at 400 MHz or 600 MHz ($^1H$), 101 MHz or 151 MHz ($^{13}C$), 377 MHz or 566 MHz ($^{19}F$) respectively. All spectra were recorded at 298 K. Chemical shifts δ are reported in ppm downfield from tetramethylsilane using the residual deuterated solvent signals as an internal reference ($CDCl_3$: $\delta_H$ = 7.26 ppm, $\delta_C$ = 77.16 ppm; $CD_3OD$: $\delta_H$ = 3.31 ppm, $\delta_C$ = 49.00 ppm; DMSO-$d_6$: $\delta_H$ = 2.50 ppm, $\delta_C$ = 39.52 ppm; $CD_3CN$: $\delta_H$ = 1.94 ppm, $\delta_C$ = 118.26 ppm). For $^1H$, $^{13}C$ and $^{19}F$ NMR, coupling constants $J$ are given in Hz and the resonance multiplicity is described as s (singlet), d (doublet), t (triplet), q (quartet), quint (quintet), sext (sextet), sept (septet), m (multiplet) and br. (broad). High-resolution mass spectrometry (HRMS) was performed by the MS-service of the EPF Lausanne (SSMI) on a Waters Xevo® G2-S Q-Tof spectrometer with electron spray ionization (ESI) or by the MS-facility of the Max Planck Institute for Medical Research on a Bruker maXis II™ ETD. Liquid chromatography coupled to mass spectrometry (LC-MS) was performed on a Shimadzu MS2020 connected to a Nexera UHPLC system equipped with a Waters ACQUITY UPLC BEH C18 (1.7 µm, 2.1 x 50 mm) column or a Supelco Titan C18 80 Å (1.9 µm, 2.1 x 50 mm). Buffer A: 0.05% HCOOH in $H_2O$ Buffer B: 0.05% HCOOH in ACN. Analytical gradient was from 10% to 90% B within 6 min with 0.5 mL/min flow unless otherwise stated. Preparative reverse phase high-performance liquid chromatography (RP-HPLC) was carried out on a Dionex system equipped with an UltiMate 3000 diode array detector for product visualization on a Waters Symmetry C18 column (5 µm, 3.9 x 150 mm), Waters SunFire™ Prep C18 OBD™ (5 µm, 10 x 150 mm) column, Supleco Ascentis® C18 column (5 µm, 10 x 250 mm) or on a Supleco Ascentis® C18 column (5 µm, 21.2 x 250 mm). Buffer A: 0.1% TFA in $H_2O$ Buffer B: ACN. Typical gradient was from 10% to 90% B within 32 min with 2, 4 or 8 mL/min flow.

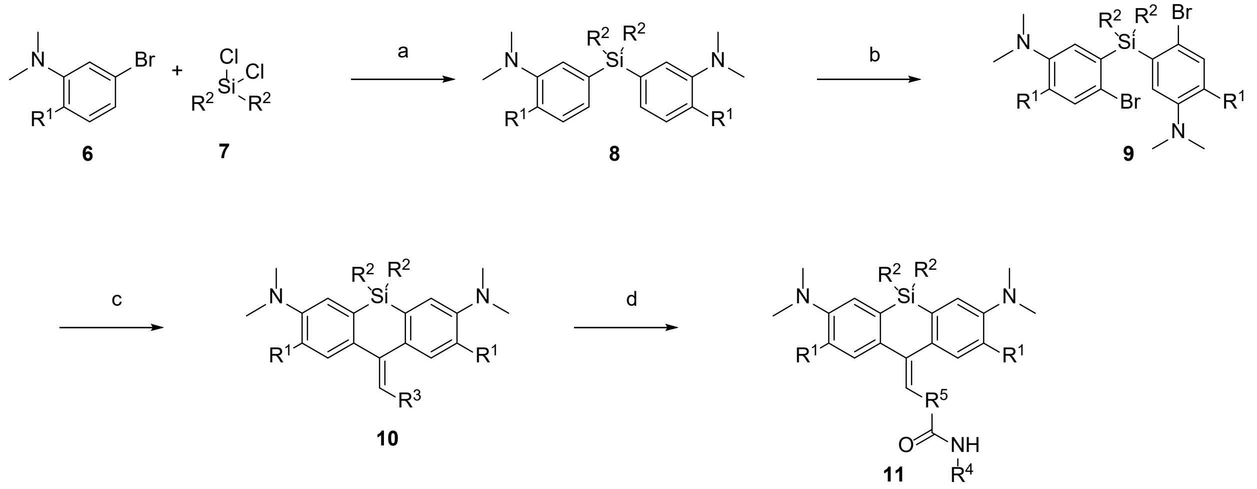


**Supplementary Scheme S1.** Synthetic route to PA-SiR derivatives according to literature procedures[3-4]: (a) *sec*-BuLi, $Et_2O$, –78 °C to room temperature 1 h; (b) NBS, $NH_4OAc$, ACN, 0 °C to room temperature, 2 h; (c) *sec*-BuLi, THF, anhydride or ester, –78 °C to room temperature 1 h; (d) TSTU, DIPEA, $R^4$-$NH_2$, DMSO, room temperature 30 min. $R^1$ = F or H; $R^2$ = Me or $^i$Pr; $R^3$ = H, $CH_2CH_3$, $CH_2COOH$ or $CH_2CH_2COOH$; $R^4$ = BG, chloroalkane, jasplakinolide; $R^5$ = $CH_2CH_2$ or $CH_2$.

General procedure **A** for the silane introduction

3-Bromo-*N,N*-dimethylaniline (**12**) (3.20 g, 16.00 mmol, 2.0 eq.) was dissolved in dry $Et_2O$ (45 mL) and cooled down to –78 °C. *sec*-BuLi (14.0 mL, 18.40 mmol, 2.3 eq., 1.3 M in cyclohexane) was added dropwise over 15 min and the mixture was stirred for 30 min at –78 °C. Dichlorodimethylsilane (**13**) (1.0 mL, 8.00 mmol, 1.0 eq.) was added dropwise over 10 min at –78 °C. The mixture was stirred for 10 min at –78 °C and then warmed up to room temperature and stirred for 1 h. The mixture was quenched with aqueous saturated $NaHCO_3$ solution. The aqueous layer was extracted with $Et_2O$ (3 x 150 mL) and the combined organic layers were dried over $MgSO_4$, filtered and evaporated to afford the crude product.

General procedure **B** for the bromination

A solution of **14** (1.85 g, 6.18 mmol, 1.0 eq.) and ammonium acetate (95 mg, 1.24 mmol, 0.2 eq.) in ACN (30 mL) was cooled down to 0 °C. NBS (2.3 g, 12.98 mmol, 2.1 eq.) was added portion wise over 10 min. The mixture was stirred at 0 °C for 30 min and then warmed up to room temperature and stirred for 2 h. A mixture of aqueous saturated $NaHCO_3$ solution and water 1:1 was added. The aqueous layer was extracted with $CH_2Cl_2$ (3 x 100 mL) and the combined organic layers were dried over $MgSO_4$, filtered and evaporated to afford the crude product.

General procedure **C** for the ring closure

A solution of **15** (365 mg, 0.8 mmol, 1.0 eq.) in dry THF (8 mL) was cooled down to –78 °C. *sec*-BuLi (1.4 mL, 1.76 mmol, 2.2 eq., 1.3 M in cyclohexane) was added dropwise over 5 min and the mixture was stirred for 30 min at –78 °C. A solution of glutaric anhydride (**16**) (100 mg, 0.88 mmol, 1.1 eq.) in dry THF (1.0 mL) was added to the mixture. The mixture was stirred at –78 °C for 15 min and then warmed up to room temperature and stirred for 30 min. Acetic acid (2 mL) was added to the mixture. The blue mixture was adsorbed on $SiO_2$ (2 g).

3,3'-(Dimethylsilanediyl)bis(*N*,*N*-dimethylaniline) **14**

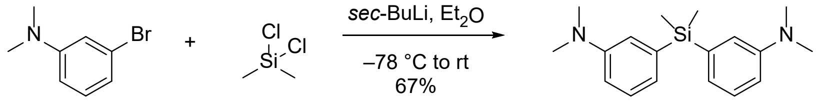


**12** **13** **14**

Following general procedure **A**, flash column chromatography ($SiO_2$, hexane/EtOAc 100:0 → 70:30) gave **14** (1.598 g, 67%) as a colorless oil.

$^1$H NMR (400 MHz, $CDCl_3$): δ 7.22 – 7.31 (m, 2H), 6.92 – 6.98 (m, 4H), 6.78 (ddd, J = 8.3, 2.8, 1.0 Hz, 2H), 2.94 (s, 12H), 0.56 (s, 6H); $^{13}$C NMR (101 MHz, $CDCl_3$): δ 150.0, 139.1, 128.6, 122.9, 118.5, 113.7, 40.8, –2.0; HRMS (*m/z*): $[M + H]^+$ calcd. for $C_{18}H_{27}N_2Si^+$, 299.1938; found, 299.1940.

3,3'-(Dimethylsilanediyl)bis(4-bromo-*N*,*N*-dimethylaniline) **15**

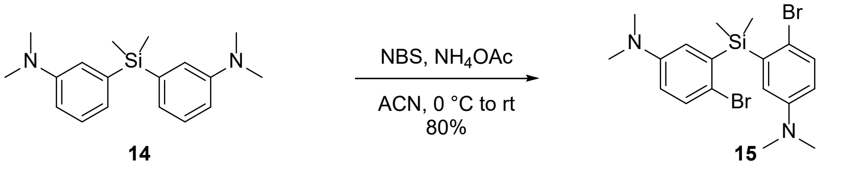


Following general procedure **B**, flash column chromatography ($SiO_2$, hexane/$CH_2Cl_2$ 100:0 → 0:100) gave **15** (2.270 g, 80%) as a beige solid.

$^1$H NMR (400 MHz, $CDCl_3$): δ 7.35 (d, J = 8.7 Hz, 2H), 6.84 (d, J = 3.2 Hz, 2H), 6.60 (dd, J = 8.7, 3.2 Hz, 2H), 2.88 (s, 12H), 0.75 (s, 6H); $^{13}$C NMR (101 MHz, $CDCl_3$): δ 149.0, 138.9, 133.1, 121.9, 116.9, 115.4, 40.7, –0.8; HRMS (m/z): $[M + H]^+$ calcd. for $C_{18}H_{25}Br_2N_2Si^+$, 455.0148; found, 455.0145.

10-(3-Carboxypropylidene)-7-(dimethylamino)-*N*,*N*,5,5-tetramethyl-5,10-dihydrodibenzo[*b*,*e*]-silin-3-aminium trifluoroacetate **PA-SiR** (**1**)

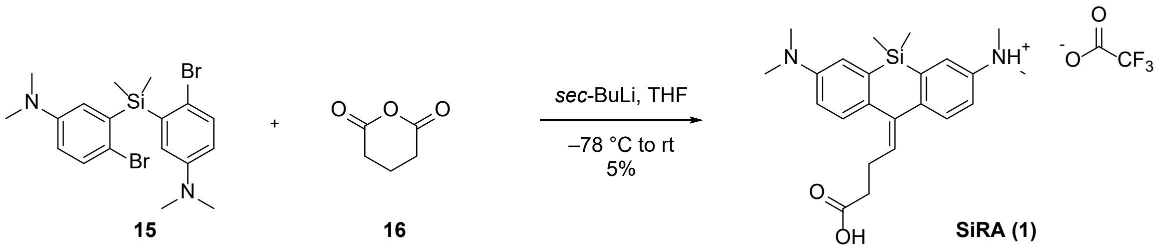


Following general procedure **C**, flash column chromatography ($SiO_2$, $CH_2Cl_2$/MeOH 100:0 → 90:10) and RP-HPLC (4 mL/min, 10% to 90% B in 32 min) gave **PA-SiR** (23 mg, 5%) as a light green solid.

$^1$H NMR (400 MHz, $CD_3OD$): δ 7.70 (t, J = 2.9 Hz, 2H), 7.66 (d, J = 8.5 Hz, 1H), 7.61 (d, J = 8.4 Hz, 1H), 7.55 (dd, J = 8.5, 2.6 Hz, 1H), 7.50 (dd, J = 8.5, 2.7 Hz, 1H), 6.03 (t, J = 7.3 Hz, 1H), 3.27 (s, 6H), 3.26 (s, 6H), 2.68 (q, J = 7.4 Hz, 2H), 2.45 (t, J = 7.2 Hz, 2H), 0.52 (br. s, 6H); $^{13}$C NMR (101 MHz, $CD_3OD$): δ 176.3, 162.3 (q, J = 36.0 Hz), 150.1, 144.5, 143.4, 143.2, 141.0, 140.3, 138.5, 134.1, 131.3, 128.6, 123.8, 123.7, 121.7, 120.0, 117.8 (q, J = 291.0 Hz), 46.4, 45.7, 34.8, 26.6, –3.8; $^{19}$F NMR (376 MHz, $CD_3OD$): δ –77.11; HRMS (*m/z*): $[M – H]^-$ calcd. for $C_{23}H_{29}N_2O_2Si^-$, 393.2004; found, 393.1992.

Note: All attempts (HPLC: Triethylammonium acetate/ACN pH = 8; triethylammonium bicarbonate/ACN pH = 8 buffer system ) to isolate the fluorescent SiR were not successful.

$N^3$,$N^3$,$N^7$,$N^7$,5,5-Hexamethyl-10-methylene-5,10-dihydrodibenzo[*b*,*e*]siline-3,7-diamine **4**

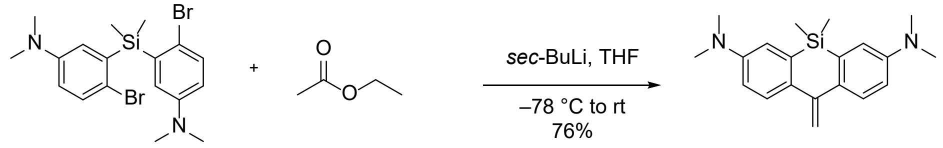


**15** **17** **4**

Following general procedure **C**, washing the solid with MeOH gave **4** (195 mg, 76%) as a white solid.

$^1$H NMR (400 MHz, $CDCl_3$): δ 7.59 (d, J = 8.7 Hz, 2H), 6.93 (d, J = 2.8 Hz, 2H), 6.79 (dd, J = 8.7, 2.8 Hz, 2H), 5.41 (s, 2H), 2.99 (s, 12H), 0.44 (s, 6H); $^{13}$C NMR (101 MHz, $CDCl_3$): δ 149.2, 147.7, 135.2, 134.8, 126.9, 115.9, 114.2, 111.3, 40.9, –2.1; HRMS (*m/z*): $[M + H]^+$ calcd. for $C_{20}H_{27}N_2Si^+$, 323.1938; found, 323.1936.

7-(Dimethylamino)-*N*,*N*,5,5-tetramethyl-10-propylidene-5,10-dihydrodibenzo[*b*,*e*]silin-3-aminium trifluoroacetate **19**

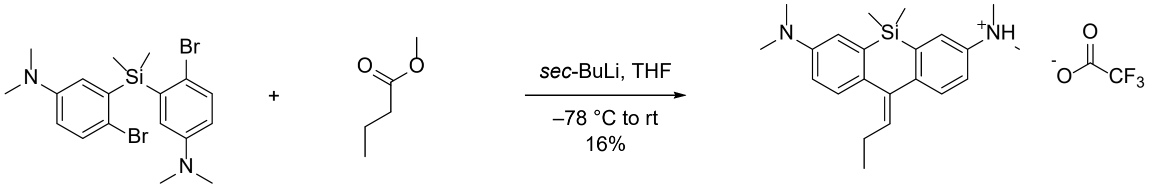


**15** **18** **19**

Following general procedure **C**, flash column chromatography ($SiO_2$, hexane/EtOAc 90:10 → 70:30) and RP-HPLC (8 mL/min, 20% to 100% B in 32 min) gave **19** (61 mg, 16%) as a light blue solid.

$^1$H NMR (400 MHz, $CD_3OD$): δ 7.60 – 7.68 (m, 2H), 7.60 (d, J = 2.7 Hz, 1H), 7.47 – 7.54 (m, 2H), 7.40 (dd, J = 8.5, 2.7 Hz, 1H), 6.01 (t, J = 7.5 Hz, 1H), 3.25 (s, 6H), 3.22 (s, 6H), 2.39 (p, J = 7.5 Hz, 2H), 1.07 (t, J = 7.5 Hz, 1H), 0.50 (br. s, 6H); $^{13}$C NMR (101 MHz, $CD_3OD$): δ 162.2 (q, J = 36.0 Hz), 150.0, 145.0, 143.6, 142.4, 139.8, 139.7, 138.3, 137.2, 131.2, 128.4, 123.3, 122.8, 121.3, 119.2, 117.7 (q, J = 283.5 Hz), 46.1, 45.2, 24.4, 14.6, –3.8; $^{19}$F NMR (376 MHz, $CD_3OD$): δ –77.15; HRMS (*m/z*): $[M + 2H]^{2+}$ calcd. for $C_{22}H_{32}N_2Si^{2+}$, 176.1162; found, 176.1160.

<u>3,7-Bis(dimethylamino)-5,5-dimethyl-3',4'-dihydro-5*H*,5'*H*-spiro[dibenzo[*b*,*e*]siline-10,2'-furan]-5'-one **21**</u>

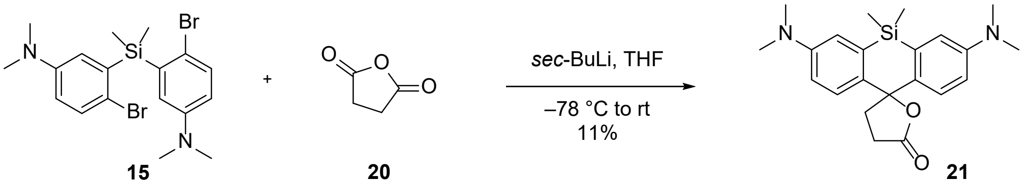


A solution of **15** (296 mg, 0.65 mmol, 1.0 eq.) in dry THF (9 mL) was cooled down to –78 °C. *sec*-BuLi (1.2 mL, 1.7 mmol, 2.6 eq., 1.3 M in cyclohexane) was added dropwise over 5 min and the mixture was stirred for 30 min at –78 °C. A solution of succinic anhydride (**20**) (72 mg, 0.72 mmol, 1.1 eq.) in dry THF (2.0 mL) was added to the mixture. The mixture was stirred at –78 °C for 15 min and then warmed up to room temperature and stirred for 30 min. Aqueous saturated $NH_4Cl$ was added and the aqueous layer was extracted with EtOAc (2 x 25 mL). The combined organic layers were dried over $MgSO_4$, filtered and evaporated to afford the crude product. Flash column chromatography ($SiO_2$, hexane/EtOAc 100:0 → 0:100) gave **21** (28 mg, 11%) as a white solid.

$^1H$ NMR (400 MHz, $CD_3CN$): δ 7.34 (d, J = 8.8 Hz, 2H), 7.06 (d, J = 2.8 Hz, 2H), 6.79 (dd, J = 8.8, 2.9 Hz, 2H), 2.95 (s, 12H), 2.52 (td, J = 7.9, 0.7 Hz, 2H), 2.30 (td, J = 8.4, 0.7 Hz, 2H), 0.58 (s, 3H), 0.42 (s, 3H); $^{13}C$ NMR (101 MHz, $CD_3CN$): δ 178.6, 150.2, 138.8, 134.8, 124.2, 118.6, 114.3, 88.7, 42.9, 40.7, 29.2, 0.5, –2.4; HRMS (m/z): $[M + H]^+$ calcd. for $C_{22}H_{29}N_2O_2Si^+$, 381.1993; found, 381.1993.

<u>10-(2-Carboxyethylidene)-7-(dimethylamino)-*N*,*N*,5,5-tetramethyl-5,10-dihydrodibenzo[b,e]-silin-3-aminium trifluoroacetate **PA-SiR-C3 (22)**</u>

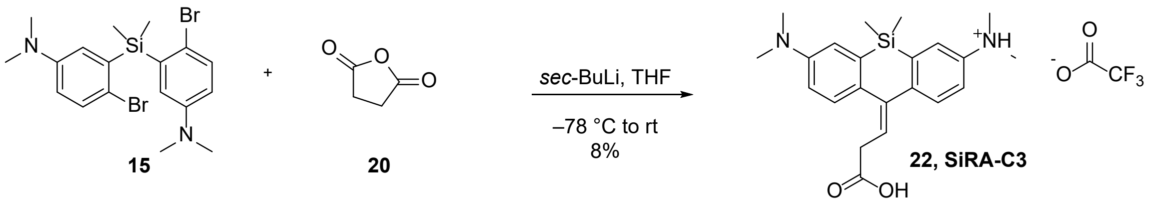


A solution of **15** (296 mg, 0.65 mmol, 1.0 eq.) in dry THF (9 mL) was cooled down to –78 °C. *sec*-BuLi (1.2 mL, 1.7 mmol, 2.6 eq., 1.3 M in cyclohexane) was added dropwise over 5 min and the mixture was stirred for 30 min at –78 °C. A solution of succinic anhydride (**20**) (72 mg, 0.72 mmol, 1.1 eq.) in dry THF (2.0 mL) was added to the mixture. The mixture was stirred at –

78 °C for 15 min and then warmed up to room temperature and stirred for 30 min. Aqueous saturated $NH_4Cl$ was added and the aqueous layer was extracted with EtOAc (2 x 25 mL). The combined organic layers were dried over $MgSO_4$, filtered and evaporated to afford the crude product. Flash column chromatography ($SiO_2$, hexane/EtOAc 100:0 → 0:100) gave **21** (28 mg, 11%) as a white solid. The compound was dissolved in water with 0.1% TFA. The blue solution was frozen and lyophilized overnight. RP-HPLC (8 mL/min, 20% to 90% B in 32 min) gave **PA-SiR-C3** (26.5 mg, 8%) as a light blue solid.

$^1$H NMR (400 MHz, $CD_3CN$): δ 7.57 – 7.60 (m, 2H), 7.48 (d, J = 2.7 Hz, 1H), 7.43 (d, J = 8.5 Hz, 1H), 7.40 (dd, J = 8.5, 2.6 Hz, 1H), 7.23 (dd, J = 8.5, 2.7 Hz, 1H), 6.06 (t, J=7.6 Hz, 1H), 3.33 (d, J = 7.7 Hz, 2H), 3.11 (s, 6H), 3.08 (s, 6H), 0.45 (s, 6H); $^{13}$C NMR (101 MHz, $CD_3CN$): δ 173.1, 147.4, 146.2, 144.4, 142.3, 139.0, 138.6, 137.5, 130.4, 127.8, 125.1, 123.1, 121.9, 120.9, 118.5, 45.3, 43.9, 35.6, -3.7; $^{19}$F NMR (376 MHz, $CD_3CN$): δ –76.26; HRMS (*m/z*): $[M + 2H]^{2+}$ calcd. for $C_{22}H_{30}N_2O_2Si^{2+}$, 191.1033; found, 191.1032.

Note: For reaction with succinic anhydride both **PA-SiR-C3** and compound **21** could be isolated. Analogues work up and isolation procedure for **PA-SiR** did not lead to the isolation of the spirolactone but only **PA-SiR**.

<u>10-(4-((2-(2-((6-Chlorohexyl)oxy)ethoxy)ethyl)amino)-4-oxobutylidene)-7-(dimethylamino)-*N*,*N*,5,5-tetramethyl-5,10-dihydrodibenzo[*b*,*e*]silin-3-aminium trifluoroacetate **PA-SiR-Halo (5)**</u>

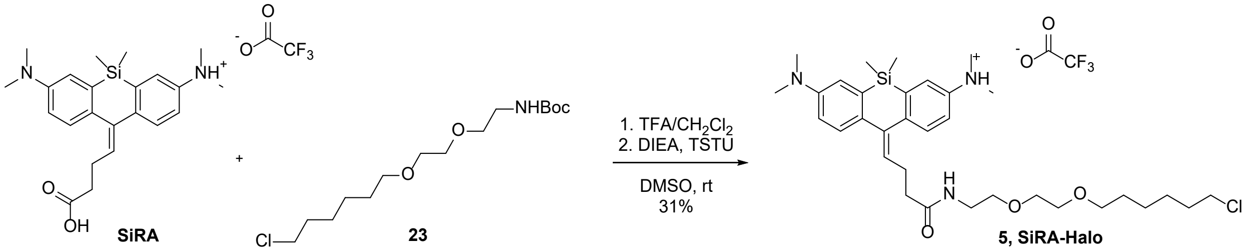


A solution of **PA-SiR** (4.0 mg, 8.0 µmol, 1.0 eq.) in DMSO (300 µL) was treated with DIEA (5.0 µL, 30.1 µmol, 3.8 eq.) and TSTU (3.6 mg, 12.0 µmol, 1.5 eq.). The mixture was shaken for 20 min at room temperature. In a separate vial a solution of Halo-NHBoc **23** (4.0 mg, 12.3 µmol, 1.5 eq.) in $TFA/CH_2Cl_2$ (2:8, 80 µL) was shaken for 5 min. The solution was evaporated and dried on the high vacuum for 1 h. The residue was taken up in DMSO (50 µL) and added to the other mixture. The mixture was shaken for 10 min and then acidified with TFA (3 µL). RP-HPLC (4 mL/min, 10% to 90% B in 32 min) gave **PA-SiR-Halo** (1.8 mg, 31%) as a light blue solid.

$^{1}$H NMR (400 MHz, $CD_3OD$): δ 7.61 (d, J = 8.5 Hz, 1H), 7.48 – 7.56 (m, 3H), 7.41 (dd, J = 8.5, 2.7 Hz, 1H), 7.30 (dd, J = 8.5, 2.7 Hz, 1H), 5.95 (t, J = 7.2 Hz, 1H), 3.48 – 3.52 (m, 8H), 3.45 (t, J = 6.5 Hz, 2H), 3.33 (d, J = 5.7 Hz, 2H), 3.22 (s, 6H), 3.19 (s, 6H), 2.69 (q, J = 7.3 Hz, 2H), 2.36 (t, J = 7.3 Hz, 2H), 1.73 (p, J = 6.9 Hz, 2H), 1.56 (p, J = 6.8 Hz, 2H), 1.30 – 1.47 (m, 4H), 0.49 (s, 6H); $^{13}$C NMR (101 MHz, $CD_3OD$): δ 175.0, 163.2 (d, J = 34.2 Hz), 148.7, 145.9, 144.4, 141.3, 139.6, 138.8, 138.2, 132.7, 131.1, 128.4, 122.5, 121.9, 120.6, 118.4, 117.7 (d, J = 291.1 Hz), 72.2, 71.2, 71.2, 70.6, 45.7, 45.6, 44.5, 40.4, 36.9, 33.7, 30.5, 27.7, 27.6, 26.5, –7.1; $^{19}$F NMR (376 MHz, $CD_3OD$): δ –77.21; HRMS (*m/z*): $[M + H]^+$ calcd. for $C_{33}H_{51}ClN_3O_3Si^+$, 600.3383; found, 600.3386.

10-(4-((4-((4*R*,7*R*,10*S*,13*S*,19*S*,*E*)-7-((1*H*-Indol-2-yl)methyl)-4-(4-hydroxyphenyl)-8,13,15,19-tetramethyl-2,6,9,12-tetraoxo-1-oxa-5,8,11-triazacyclononadec-15-en-10-yl)butyl)amino)-4-oxobutylidene)-7-(dimethylamino)-*N*,*N*,5,5-tetramethyl-5,10-dihydrodibenzo[*b*,*e*]silin-3-aminium trifluoroacetate **PA-SiR-Actin (25)**

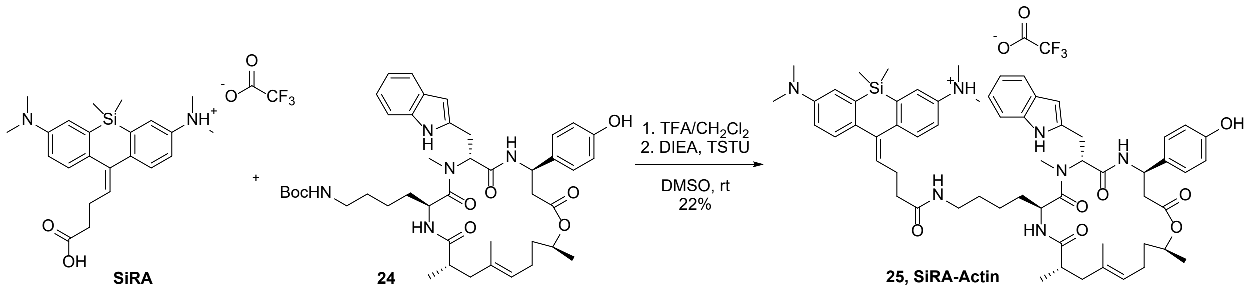


A solution of **PA-SiR** (4.0 mg, 8.0 µmol, 1.0 eq.) in DMSO (300 µL) was treated with DIEA (5.0 µL, 30.1 µmol, 3.8 eq.) and TSTU (3.6 mg, 12.0 µmol, 1.5 eq.). The mixture was shaken for 20 min at room temperature. In a separate vial a solution of jasplakinolide-NHBoc **24** (9.5 mg, 12.3 µmol, 1.5 eq.) in TFA/$CH_2Cl_2$ (2:8, 80 µL) was shaken for 2 min. The solution was evaporated and dried on the high vacuum for 1 h. The residue was taken up in DMSO (50 µL) and added to the other mixture. The mixture was shaken for 30 min and then acidified with TFA (3 µL). RP-HPLC (4 mL/min, 10% to 90% B in 32 min) gave **PA-SiR-Actin** (2.6 mg, 22%) as a light blue solid.

$^{1}$H NMR (400 MHz, $CD_3OD$): δ 8.42 (d, J = 8.6 Hz, 1H), 7.64 (d, J = 8.5 Hz, 1H), 7.53 – 7.61 (m, 4H), 7.47 (dd, J = 8.4, 2.7 Hz, 1H), 7.32 – 7.37 (m, 1H), 7.24 – 7.30 (m, 1H), 6.95 – 7.10 (m, 5H), 6.72 – 6.81 (m, 2H), 5.96 (t, J = 7.3 Hz, 1H), 5.60 (dd, J = 10.1, 6.4 Hz, 1H), 5.27 (dt, J = 9.8, 5.0 Hz, 1H), 5.04 (t, J = 7.0 Hz, 1H), 4.78 – 4.85 (m, 1H), 4.67 (t, J = 5.7 Hz, 1H), 3.23 (s, 6H), 3.18

(s, 6H), 2.99 – 3.15 (m, 4H), 2.93 (s, 3H), 2.66 – 2.79 (m, 4H), 2.61 (ddd, J = 10.2, 6.8, 3.0 Hz, 1H), 2.35 (t, J = 7.2 Hz, 2H), 2.26 – 2.33 (m, 1H), 1.84 – 1.95 (m, 3H), 1.57 – 1.64 (m, 1H), 1.55 (s, 3H), 1.41 (td, J = 13.3, 7.7 Hz, 1H), 1.23 (s, 2H), 1.18 (d, J = 6.3 Hz, 3H), 1.07 (d, J = 6.8 Hz, 3H), 0.95 (d, J = 5.5 Hz, 2H), 0.86 (d, J = 8.1 Hz, 2H), 0.53 (d, J = 3.6 Hz, 6H); $^{13}C$ NMR (101 MHz, $CD_3OD$): δ 178.0, 174.9, 174.4, 172.3, 171.7, 157.8, 149.0, 145.8, 144.2, 141.3, 139.6, 138.3, 138.0, 135.4, 134.7, 133.6, 132.9, 131.1, 128.5, 128.4, 128.2, 125.8, 124.4, 122.9, 122.4, 120.9, 119.8, 119.5, 119.0, 118.6, 116.3, 112.3, 110.7, 71.9, 57.4, 50.7, 50.1, 45.7, 44.6, 44.5, 41.9, 40.3, 40.1, 37.0, 36.6, 32.6, 31.3, 30.0, 27.5, 25.9, 24.6, 23.2, 20.3, 20.0, 16.6, –0.2; $^{19}F$ NMR (376 MHz, $CD_3OD$): δ –77.26; HRMS (*m/z*): $[M + H]^+$ calcd for $C_{61}H_{80}N_7O_7Si^+$, 1050.5883; found, 1050.5902.

10-(4-((4-(((2-Amino-9*H*-purin-6-yl)oxy)methyl)benzyl)amino)-4-oxobutylidene)-7-(dimethylamino)-*N*,*N*,5,5-tetramethyl-5,10-dihydrodibenzo[*b*,*e*]silin-3-aminium trifluoroacetate **PA-SiR-SNAP (27)**

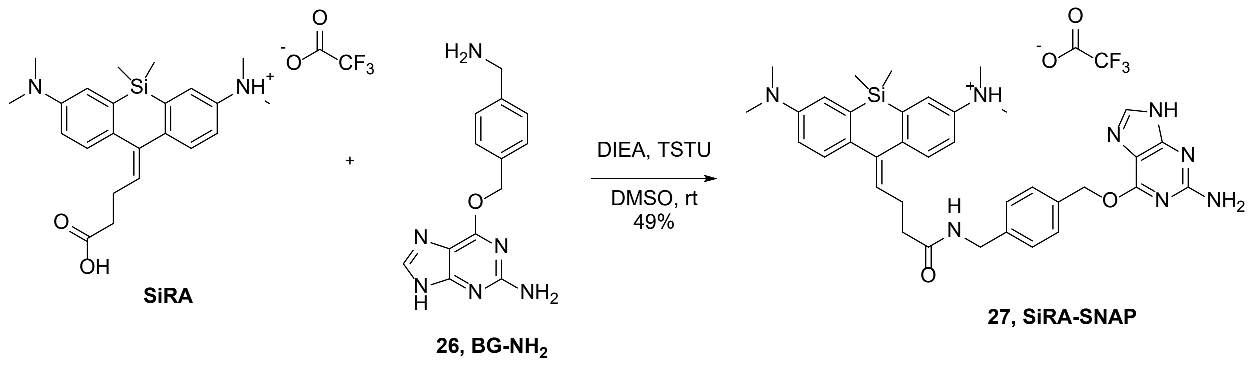


A solution of **PA-SiR** (4.0 mg, 8.0 µmol, 1.0 eq.) in DMSO (300 µL) was treated with DIEA (5.0 µL, 30.1 µmol, 3.8 eq.) and TSTU (3.6 mg, 12.0 µmol, 1.5 eq.). The mixture was shaken for 20 min and BG-$NH_2$ **26** (3.4 mg, 12.8 µmol, 1.6 eq.) was added. The mixture was shaken for 10 min at room temperature and then acidified with TFA (15 µL). RP-HPLC (8 mL/min, 10% to 90% B in 32 min) gave **PA-SiR-SNAP** (3.0 mg, 49%) as a light green solid.

$^1H$ NMR (400 MHz, $CD_3OD$): δ 8.32 (s, 1H), 7.47 – 7.58 (m, 4H), 7.36 – 7.42 (m, 3H), 7.25 – 7.30 (m, 3H), 5.95 (t, J = 7.2 Hz, 1H), 5.60 (s, 2H), 4.35 (s, 2H), 3.21 (s, 6H), 3.17 (s, 6H), 2.73 (q, J = 7.2 Hz, 2H), 2.41 (t, J = 7.2 Hz, 2H), 0.46 (s, 6H); $^{13}C$ NMR (101 MHz, $CD_3OD$): δ 174.9, 162.2 (d, J = 36.2 Hz), 161.1, 158.6, 153.8, 148.3, 146.1, 144.6, 143.2, 141.5, 140.8, 140.4, 139.6, 138.1, 135.4, 132.4, 131.1, 130.1, 128.7, 128.3, 122.4, 121.8, 120.5, 118.2, 117.8 (d, J = 290.6

Hz), 108.7, 70.6, 45.4, 44.3, 43.7, 37.0, 27.6, –2.6; $^{19}$F NMR (376 MHz, $CD_3OD$): δ –77.14; HRMS (*m/z*): $[M + H]^+$ calcd. for $C_{37}H_{43}N_8O_2Si^+$, 647.3278; found, 647.3274.

<u>10-(3-((2-(2-((6-Chlorohexyl)oxy)ethoxy)ethyl)amino)-3-oxopropylidene)-7-(dimethylamino)-*N*,*N*,5,5-tetramethyl-5,10-dihydrodibenzo[*b*,*e*]silin-3-aminium trifluoroacetate **PA-SiR-C3-Halo (28)**</u>

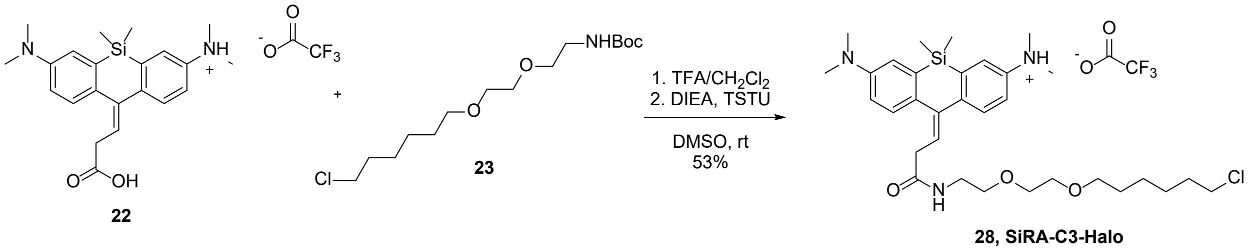


A solution of **22** (4.0 mg, 8.1 µmol, 1.0 eq.) in DMSO (250 µL) was treated with DIEA (5.2 µL, 31.5 µmol, 4.0 eq.) and TSTU (3.8 mg, 12.6 µmol, 1.6 eq.). The mixture was shaken for 20 min at room temperature. In a separate vial a solution of Halo-NHBoc **23** (4.8 mg, 14.7 µmol, 1.8 eq.) in TFA/$CH_2Cl_2$ (2:8, 200 µL) was shaken for 5 min. The solution was evaporated and dried on the high vacuum for 1 h. The residue was taken up in DMSO (150 µL) and added to the other mixture. The mixture was shaken for 10 min and then acidified with TFA (15 µL). RP-HPLC (4 mL/min, 20% to 100% B in 32 min) gave **PA-SiR-C3-Halo** (3.0 mg, 53%) as a light blue solid.

$^1$H NMR (400 MHz, $CD_3OD$): δ 7.70 (d, *J* = 8.5 Hz, 1H), 7.59 (d, *J* = 2.6 Hz, 1H), 7.55 (d, *J* = 8.5 Hz, 1H), 7.50 – 7.41 (m, 2H), 7.24 (dd, *J* = 8.5, 2.7 Hz, 1H), 6.13 (t, *J* = 7.6 Hz, 1H), 3.64 – 3.48 (m, 6H), 3.52 (t, J = 6.6 Hz, 2H), 3.46 (t, J = 6.5 Hz, 2H), 3.39 (t, J = 5.4 Hz, 2H), 3.30 (d, J = 3.3 Hz, 2H), 3.24 (s, 6H), 3.16 (s, 6H), 1.72 (dq, J = 7.9, 6.7 Hz, 2H), 1.55 (tt, J = 7.5, 6.4 Hz, 2H), 1.49 – 1.22 (m, 4H), 0.49 (s, 6H); $^{13}$C NMR (101 MHz, $CD_3OD$): δ 173.7, 161.9 (q, J = 36.5 Hz), 149.0, 147.0, 144.2, 143.2, 139.3, 138.7, 138.5, 131.0, 128.5, 126.3, 122.8, 121.2, 120.8, 117.7, 117.7 (q, J = 290.2 Hz), 72.2, 71.2, 71.2, 70.5, 45.8, 45.7, 43.8, 40.5, 38.1, 33.7, 30.5, 27.7, 26.4, –3.5; $^{19}$F NMR (376 MHz, $CD_3OD$): δ –77.2; HRMS (m/z): $[M + H]^+$ calcd. for $C_{32}H_{49}ClN_3O_3Si^+$, 586.3226; found, 586.3226.

3,3'-(Diisopropylsilanediyl)bis(*N*,*N*-dimethylaniline) **30**

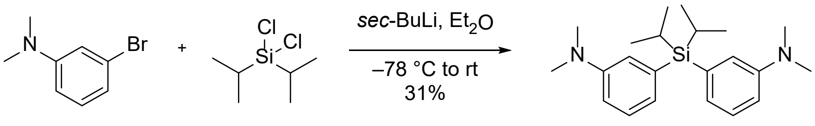


12 29 30

Following general procedure **A**, flash column chromatography ($SiO_2$, hexane/EtOAc 100:0 → 80:20) gave **30** (0.440 g, 31%) as a colorless oil.

$^{1}$H NMR (400 MHz, $CDCl_3$): δ 7.21 – 7.29 (m, 2H), 6.90 – 7.00 (m, 4H), 6.81 (ddd, J = 8.3, 2.7, 1.0 Hz, 2H), 2.92 (s, 12H), 1.56 (hept, J = 7.3 Hz, 2H), 0.97 (d, J = 7.4 Hz, 12H); $^{13}$C NMR (101 MHz, $CDCl_3$): δ 149.7, 133.9, 128.1, 125.2, 120.9, 113.7, 41.0, 17.8, 10.0; HRMS (*m/z*): $[M + H]^+$ calcd. for $C_{22}H_{35}N_2Si^+$, 355.2564; found, 355.2567.

3,3'-(Diisopropylsilanediyl)bis(4-bromo-*N*,*N*-dimethylaniline) **31**

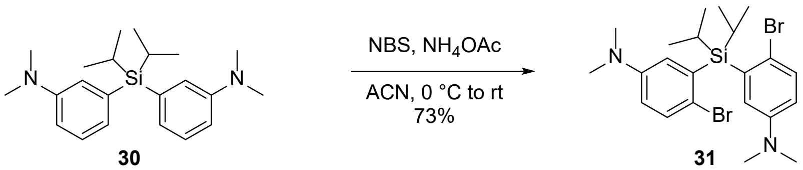


30 31

Following general procedure **B**, flash column chromatography ($SiO_2$, hexane/$CH_2Cl_2$ 100:0 → 0:100) gave **31** (1.087 g, 73%) as a white solid.

$^{1}$H NMR (400 MHz, $CDCl_3$): δ 7.33 (d, J = 8.7 Hz, 2H), 6.97 (d, J = 3.2 Hz, 2H), 6.60 (dd, J = 8.8, 3.3 Hz, 2H), 2.92 (s, 12H), 2.00 (p, J = 7.4 Hz, 2H), 1.17 (d, J = 7.5 Hz, 12H); $^{13}$C NMR (101 MHz, $CDCl_3$): δ 148.6, 136.8, 133.4, 122.8, 117.6, 115.1, 40.8, 19.1, 12.7; HRMS (*m/z*): $[M + H]^+$ calcd. for $C_{22}H_{33}Br_2N_2Si^+$, 511.0774; found, 511.0779.

10-(3-Carboxypropylidene)-7-(dimethylamino)-5,5-diisopropyl-*N*,*N*-dimethyl-5,10-dihydrodibenzo[*b*,*e*]silin-3-aminium trifluoroacetate **32**

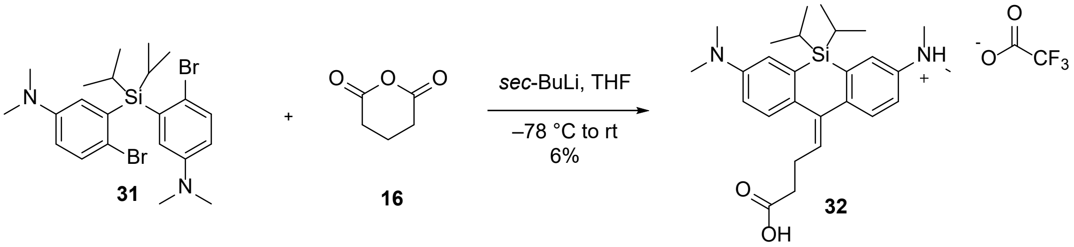


31 16 32

Following general procedure **C**, flash column chromatography ($SiO_2$, hexane/EtOAc 100:0 → 50:50) and RP-HPLC (2 mL/min, 10% to 90% B in 32 min) gave **32** (6.7 mg, 6%) as a light blue solid.

$^1H$ NMR (400 MHz, $CD_3OD$): δ 7.69 (d, J = 8.5 Hz, 1H), 7.48 – 7.61 (m, 4H), 7.37 (dd, J = 8.6, 2.7 Hz, 1H), 5.90 (t, J = 7.2 Hz, 1H), 3.24 (s, 6H), 3.20 (s, 6H), 2.65 (q, J = 7.6 Hz, 2H), 2.41 (t, J = 7.4 Hz, 2H), 1.57 (br. s, 2H), 1.09 (br. s, 12H); $^{13}C$ NMR (101 MHz, $CD_3OD$): δ 176.5, 161.9 (q, J = 36.2 Hz), 150.5, 145.7, 143.9, 142.2, 141.7, 136.0, 134.7, 133.7, 131.6, 128.7, 123.9, 123.0, 121.0, 116.3 (q, J = 292.8 Hz), 45.8, 44.5, 34.8, 26.9, 18.4, 12.9; $^{19}F$ NMR (376 MHz, $CD_3OD$): δ –77.16; HRMS (*m/z*): $[M – H]^–$ calcd. for $C_{27}H_{37}N_2O_2Si^–$, 449.2630; found, 449.2622.

5-Bromo-2-fluoro-*N*,*N*-dimethylaniline **34**

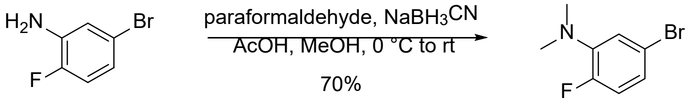


**33** **34**

A solution of 5-bromo-2-fluoroaniline **33** (5.0 g, 26.3 mmol, 1.0 eq.) in MeOH (30 mL) was treated with acetic acid (40 mL) and paraformaldehyde (3.9 g, 131.6 mmol, 5.0 eq.). The mixture was cooled down to 0 °C and stirred for 15 min. $NaBH_3CN$ (5.0 g, 78.9 mmol, 3.0 eq.) was added portion wise to the mixture over 10 min. The mixture was warmed up to room temperature and was stirred for 16 h. The mixture was evaporated and then neutralized with aqueous NaOH solution (4 mL, 5 M). The aqueous layer was extracted with $CH_2Cl_2$ (3 x 100 mL). The combined organic layers were dried over $MgSO_4$, filtered and evaporated to afford the crude product. Flash column chromatography ($SiO_2$, hexane/EtOAc 100:0 → 85:15) gave **34** (3.990 g, 70%) as a yellow oil.

$^1H$ NMR (400 MHz, $CDCl_3$): δ 6.82 – 6.99 (m, 3H), 2.85 (d, J = 1.0 Hz, 6H); $^{13}C$ NMR (101 MHz, $CDCl_3$): δ 154.1 (d, J = 245.3 Hz), 142.1 (d, J = 9.7 Hz), 123.3 (d, J = 7.9 Hz), 121.2 (d, J = 3.9 Hz), 117.6 (d, J = 22.8 Hz), 116.9 (d, J = 3.3 Hz), 42.7 (d, J = 4.5 Hz); $^{19}F$ NMR (376 MHz, $CDCl_3$): δ –124.69 – –124.58 (m); HRMS (*m/z*): $[M + H]^+$ calcd. for $C_8H_{10}BrFN^+$, 217.9975; found, 217.9978.

5,5'-(Dimethylsilanediyl)bis(2-fluoro-*N*,*N*-dimethylaniline) **35**

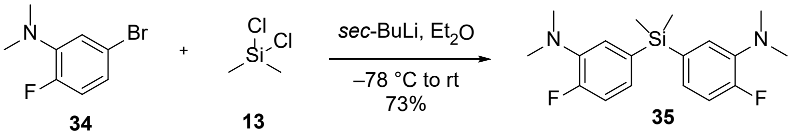


Following general procedure **A**, flash column chromatography ($SiO_2$, hexane/EtOAc 100:0 → 85:15) gave **35** (2.110 g, 73%) as a yellow oil.

$^1$H NMR (400 MHz, $CDCl_3$): δ 6.98 – 7.06 (m, 6H), 2.83 (d, J = 0.9 Hz, 12H), 0.52 (s, 6H); $^{13}$C NMR (101 MHz, $CDCl_3$): δ 156.3 (d, J = 248.0 Hz), 140.3 (d, J = 8.0 Hz), 134.0 (d, J = 4.3 Hz), 127.6 (d, J = 7.5 Hz), 124.0 (d, J = 3.4 Hz), 115.9 (d, J = 19.8 Hz), 43.0 (d, J = 3.9 Hz), –1.9; $^{19}$F NMR (376 MHz, $CDCl_3$): δ –121.34 – –121.21 (m); HRMS (*m/z*): $[M + H]^+$ calcd. for $C_{18}H_{25}F_2N_2Si^+$, 335.1750; found, 335.1754.

5,5'-(Dimethylsilanediyl)bis(4-bromo-2-fluoro-*N*,*N*-dimethylaniline) **36**

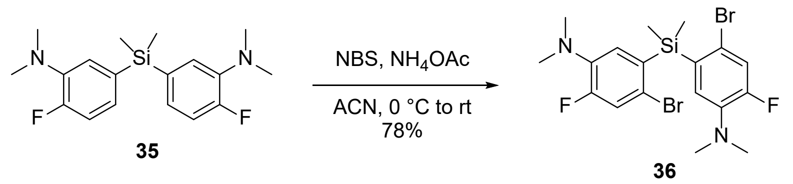


Following general procedure **B**, flash column chromatography ($SiO_2$, hexane/$CH_2Cl_2$ 100:0 → 0:100) gave **36** (2.329 g, 78%) as a beige solid.

$^1$H NMR (400 MHz, $CDCl_3$): δ 7.20 (d, J = 12.5 Hz, 2H), 6.94 (d, J = 10.2 Hz, 2H), 2.80 (d, J = 1.0 Hz, 12H), 0.74 (s, 6H); $^{13}$C NMR (101 MHz, $CDCl_3$): δ 155.6 (d, J = 252.7 Hz), 139.4 (d, J = 7.5 Hz), 134.1 (d, J = 4.0 Hz), 126.7 (d, J = 4.0 Hz), 120.9 (d, J = 23.2 Hz), 119.9 (d, J = 8.6 Hz), 42.7 (d, J = 4.0 Hz), –0.8; $^{19}$F NMR (376 MHz, $CDCl_3$): δ –118.89 (t, J = 11.3 Hz); HRMS (*m/z*): $[M + H]^+$ calcd. for $C_{18}H_{23}Br_2F_2N_2Si^+$, 490.9960; found, 490.9954.

10-(3-Carboxypropylidene)-7-(dimethylamino)-2,8-difluoro-*N*,*N*,5,5-tetramethyl-5,10-dihydrodibenzo[*b*,*e*]silin-3-aminium trifluoroacetate **37**

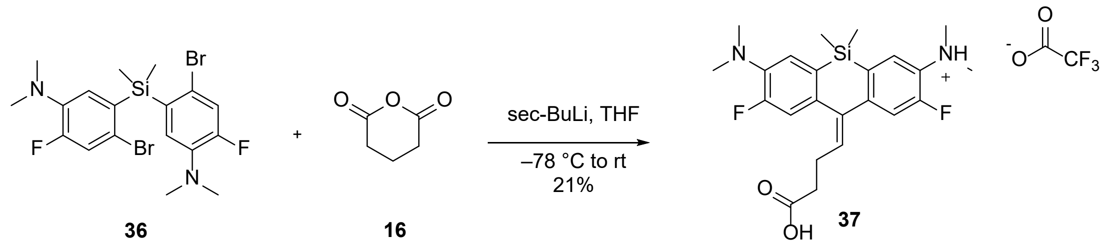


Following general procedure **C**, flash column chromatography ($SiO_2$, hexane/EtOAc 80:20 → 0:100) and RP-HPLC (3 mL/min, 10% to 90% B in 32 min) gave **37** (24 mg, 21%) as a white solid.

$^1$H NMR (400 MHz, $CD_3OD$): δ 7.58 (d, J = 9.0 Hz, 1H), 7.54 (d, J = 9.3 Hz, 1H), 7.40 (d, J = 13.3 Hz, 1H), 7.35 (d, J = 13.6 Hz, 1H), 6.00 (t, J = 7.3 Hz, 1H), 3.15 (s, 6H), 3.10 (s, 6H), 2.69 (q, J = 7.2 Hz, 2H), 2.47 (t, J = 7.1 Hz, 2H), 0.48 (s, 6H); $^{13}$C NMR (101 MHz, $CD_3OD$): δ 176.4, 163.2 (d, J = 34.8 Hz), 161.1 (q, J = 38.1 Hz), 157.8 (d, J = 85.7 Hz), 155.3 (d, J = 84.7 Hz), 150.0 (d, J = 7.1 Hz), 142.5 (d, J = 5.5 Hz), 139.9, 135.7 (d, J = 8.2 Hz), 134.9 (d, J = 3.5 Hz), 134.5, 133.7 (d, J = 11.7 Hz), 133.3 (d, J = 3.9 Hz), 124.9, 118.0 (d, J = 20.3 Hz), 117.2 (q, J = 288.5 Hz), 115.3 (d, J = 20.0 Hz), 45.3 (d, J = 2.5 Hz), 44.7 (d, J = 2.9 Hz), 34.7, 26.5, –3.4; $^{19}$F NMR (376 MHz, $CD_3OD$): δ –77.45, –123.44 (dd, J = 13.3, 9.3 Hz); HRMS (*m/z*): $[M-H]^-$ calcd. for $C_{23}H_{27}F_2N_2O_2Si^-$, 429.1815; found, 429.1816.

*General Considerations for In Vitro Characterization and Microscopy*

**PA-SiRs** were prepared as stock solutions in dry DMSO and diluted in the respective buffer such that the final concentration of DMSO did not exceed 5% v/v. A fiber coupled LED (Omicron, 340 nm, 3 mm liquid light guide) was used to perform UV irradiation unless otherwise stated. PBS (6.7 mM, Lonza) was used in all experiments.

*UV-Vis Measurements*

All absorbance measurements (spectra and time traces) were performed in 1.5 mL stirrable quartz cuvettes (Hellma Analytics) on a JASCO V770 spectrophotometer with a Peltier element (PAC743R) under continuous stirring and at 21 °C **PA-SiRs** were diluted in PBS (10 µM unless otherwise stated). A blank was measured before starting the measurement. UV irradiation was performed directly inside the spectrophotometer during the ongoing experiment for 12 s unless otherwise stated. PBS solutions of different pH were adjusted by addition of HCl or NaOH solution using a pH meter. Cysteamine concentrations were adjusted by the addition of concentrated cysteamine solution (1 M). **PA-SiR-Halo**, **PA-SiR-SNAP** and **PA-SiR-Actin** probes (10 µM) were directly added to the target protein (20 µM SNAP-tag, 20 µM HaloTag or 0.4 mg/mL G-actin), or to a bovine serum albumin (BSA) (Sigma) solution in PBS. The mixture was incubated for 1 h (HaloTag) or 2 h (SNAP-tag) at room temperature. In the case of the actin probe, buffer containing 5 mM Tris-HCl (pH 8.0), 0.2 mM $CaCl_2$ and 0.2 mM ATP was used. This buffer was supplemented with 50 mM KCl, 2 mM $MgCl_2$, 5 mM guanidine carbonate and 1 mM ATP to obtain F-actin. Both buffers are components of the actin polymerization fluorescence assay kit (Cytoskeleton). The samples were incubated for 2–3 h at 37 °C. Measurements were performed in triplicates. Fitted parameters such as decay constants etc. are reported as the average of three fits. Representative measurements are displayed. Where given $\boldsymbol{X^2}$ the reduced chi-squared corresponds to the residual sum of square (RSS) and $\boldsymbol{R^2}$ is the squared correlation coefficient. They are defined as follows:

$$X^2 = RSS = \sum_{i=1}^{n} (y_i - \hat{y_i})^2$$

$$R^2 = 1 - \frac{RSS}{TSS} = 1 - \frac{\sum_{i=1}^{n}(y_i - \hat{y_i})^2}{\sum_{i=1}^{n}(y_i - \bar{y_i})^2}$$

*Fluorescence Measurements and Determination of Quantum Yield*

Fluorescence spectra were measured on a JASCO FP-8600 fluorimeter in 1.4 mL fluorescence cuvettes (Hellma Analytics). Emission spectra were collected from 610–1000 nm exciting at 580 nm; excitation spectra were recorded at 664 nm exciting from 400–655 nm unless otherwise stated. Fluorescence intensity upon addition of cysteamine was measured on a plate reader (TECAN Spark® 20M) equipped with a monochromator exciting at 640/10 nm and collecting the emission at 670/10 nm. Quantum yields were determined using a Hamamatsu Quantaurus QY.

*Determination of Extinction Coefficient at 646 nm and Quantum Yield of Activation*

Extinction coefficients at 646 nm after activation were calculated from the equilibrium constants ($K_2$) obtained in the 12 s activation experiments (Fig. 2b, Supplementary Fig. S4, S7, Table S4 and S6), assuming that during the activation the decay ($k_2$ and $k_{-2}$) is negligible, and the absorbance reached in equilibrium in the saturation experiment (Supplementary Fig. S8, Table S7). Quantum yields of activation were determined using standard ferroxialate actinometry[5] along with the activation rates determined in the saturation experiments (Supplementary Fig. S8). This calculation does not take into account the decay kinetics but was good enough to give an estimate of the quantum yields of activation.

*Computational Chemistry*

Optimization of the **PA-SiR** structure as well as HOMO/LUMO calculations were performed at the B3LYP/6-31G(d) level of theory by using the software package Gaussian 09[6].

*LC-MS Analysis*

LC-MS was performed on a Shimadzu MS2020 connected to a Nexera UHPLC system equipped with a Supelco Titan C18 80 Å (1.9 µm, 2.1 x 50 mm) column. Buffer A: 0.05% HCOOH in $H_2O$ Buffer B: 0.05% HCOOH in ACN. **PA-SiR** was dissolved in MQ water (~20 µM). UV irradiation was performed for 1 min in a quartz cuvette (Hellma Analytics) and aliquots were taken to measure LC-MS at defined time points using an analytical gradient from 10% to 90% B within 6 min with 0.5 mL/min flow.

*$^1$H NMR Analysis*

**PA-SiR** (1 mg, 2.0 μmol) was dissolved in PBS/$D_2O$ (1 mL, 90:10) and NaOH (1 μL, 5 M) was added to achieve better solubility as **PA-SiR** was isolated as its TFA salt (pH = 7–8, pH paper). $^1$H NMR spectra were measured on a Bruker AV 600 spectrometer at 600 MHz and 298 K. Chemical shifts δ are reported in ppm downfield from tetramethylsilane using the DMSO signal ($\delta_H$ = 2.50 ppm) instead of the residual deuterated solvent signal as an internal reference. Spectra were measured with NS = 128 using a water suppression pre-saturation sequence. UV irradiation was performed outside of the spectrometer for the indicated times with a transilluminator (Biometra TI 1, 312 nm). After each UV irradiation step the NMR sample was transferred to the NMR spectrometer.

*Plasmids*

A pcDNA5/FRT/TO vector (ThermoFisher Scientific) was used for transient expression in mammalian cells and generation of stable cell lines. A pET51b(+) vector (Novagen) was used for protein production in *Escherichia coli*. Proteins were tagged Strep and $His_{x10}$ N- and C-terminal, respectively. SNAP-tag and HaloTag7 were fused to the N or C terminus of the genes of interest (GOI) and a T2A-EGFP sequence was introduced. Cloning was performed by Gibson assembly[7]. GOI: H2B (NEB, pSNAPf-H2B), CEP41 (Genecopoeia (GC-V1653 and GC-V1653-CF)),[8] mEOS3.2 (Addgene #54525)[9], Lifeact (Addgene #36201)[10], TOMM20 (Addgene #55146, gift from Michael Davidson), β-2-adrenergic-receptor-Halo (Addgene #66994, gift from Catherine Berlot) were used as entry plasmids.

*Protein Production and Purification*

Proteins were expressed in *E. coli* strain BL21(DE3)-pLysS. Luria-Bertani broth (LB) cultures were grown at 37 °C to optical density at 600 nm ($OD_{600nm}$) of 0.8, induced by the addition of 0.5 mM isopropyl-β-D-thiogalactopyranoside and grown at 17 °C overnight in the presence of 1 mM $MgCl_2$. The cells were harvested by centrifugation (4'500 g, 10 min, 4 °C) and lysed by sonication. The cell lysate was cleared by centrifugation (20'000 g, 20 min, 4 °C). All proteins were purified using affinity-tag Ni-NTA (Qiagen) leading to higher than 95% pure proteins (verified by SDS-PAGE coomassie staining). mEOS3.2-Halo was purified analogously but using an additional Strep-Tactin (IBA) column purification step to reach higher purity and following the suppliers' instructions. Proteins were finally concentrated using an Ultra-0.5 mL centrifugal filter device (Amicon) with a molecular weight cut-off (MWCO) according to the protein size and then stored in a glycerol 45% solution at –20 °C. Proteins were used from glycerol stocks and were further diluted.

*Protein amino acid sequences*

HaloTag

MASWSHPQFEKGADDDDKVPHGSEIGTGFPFDPHYVEVLGERMHYVDVGPRDGTPVLF
LHGNPTSSYVWRNIIPHVAPTHRCIAPDLIGMGKSDKPDLGYFFDDHVRFMDAFIEALGL
EEVVLVIHDWGSALGFHWAKRNPERVKGIAFMEFIRPIPTWDEWPEFARETFQAFRTTD
VGRKLIIDQNVFIEGTLPMGVVRPLTEVEMDHYREPFLNPVDREPLWRFPNELPIAGEPA
NIVALVEEYMDWLHQSPVPKLLFWGTPGVLIPPAEAARLAKSLPNCKAVDIGPGLNLLQ
EDNPDLIGSEIARWLSTLEISGAPGFSSISAHHHHHHHHHH

Green: Strep-tag, Red: HaloTag, Purple: His-tag

SNAP-tag

MASWSHPQFEKGADDDDKVPHMDKDCEMKRTTLDSPLGKLELSGCEQGLHEIIFLGKG
TSAADAVEVPAPAAVLGGPEPLMQATAWLNAYFHQPEAIEEFPVPALHHPVFQQESFTR
QVLWKLLKVVKFGEVISYSHLAALAGNPAATAAVKTALSGNPVPILIPCHRVVQGDLDV
GGYEGGLAVKEWLLAHEGHRLGKPGLGAPGFSSISAHHHHHHHHHH

Green: Strep-tag, Red: SNAP-tag, Purple: His-tag

mEOS3.2:HaloTag

MASWSHPQFEKGADDDDKVPHMSAIKPDMKIKLRMEGNVNGHHFVIDGDGTGKPFEG
KQSMDLEVKEGGPLPFAFDILTTAFHYGNRVFAKYPDNIQDYFKQSFPKGYSWERSLTF
EDGGICNARNDITMEGDTFYNKVRFYGTNFPANGPVMQKKTLKWEPSTEKMYVRDGV
LTGDIEMALLLEGNAHYRCDFRTTYKAKEKGVKLPGAHFVDHCIEILSHDKDYNKVKL
YEHAVAHSGLPDNARRGRLEVLFQGPKAFLEGSEIGTGFPFDPHYVEVLGERMHYVDV
GPRDGTPVLFLHGNPTSSYVWRNIIPHVAPTHRCIAPDLIGMGKSDKPDLGYFFDDHVRF
MDAFIEALGLEEVVLVIHDWGSALGFHWAKRNPERVKGIAFMEFIRPIPTWDEWPEFAR
ETFQAFRTTDVGRKLIIDQNVFIEGTLPMGVVRPLTEVEMDHYREPFLNPVDREPLWRFP

NELPIAGEPANIVALVEEYMDWLHQSPVPKLLFWGTPGVLIPPAEAARLAKSLPNCKAV
DIGPGLNLLQEDNPDLIGSEIARWLSTLEISGAPGFSSISAHHHHHHHHHH
Green: Strep-tag, Red: HaloTag, Blue: mEOS3.2, Purple: His-tag

SNAP-tag:EGFP:HaloTag
MASWSHPQFEKGADDDDKVPHMDKDCEMKRTTLDSPLGKLELSGCEQGLHEIIFLGKG
TSAADAVEVPAPAAVLGGPEPLMQATAWLNAYFHQPEAIEEFPVPALHHPVFQQESFTR
QVLWKLLKVVKFGEVISYSHLAALAGNPAATAAVKTALSGNPVPILIPCHRVVQGDLDV
GGYEGGLAVKEWLLAHEGHRLGKPGLGGRLEVLFQGPKAFLEMVSKGEELFTGVVPIL
VELDGDVNGHKFSVSGEGEGDATYGKLTLKFICTTGKLPVPWPTLVTTLTYGVQCFSRY
PDHMKQHDFFKSAMPEGYVQERTIFFKDDGNYKTRAEVKFEGDTLVNRIELKGIDFKED
GNILGHKLEYNYNSHNVYIMADKQKNGIKVNFKIRHNIEDGSVQLADHYQQNTPIGDGP
VLLPDNHYLSTQSALSKDPNEKRDHMVLLEFVTAAGITLGMDELYKIGTGFPFDPHYVE
VLGERMHYVDVGPRDGTPVLFLHGNPTSSYVWRNIIPHVAPTHRCIAPDLIGMGKSDKP
DLGYFFDDHVRFMDAFIEALGLEEVVLVIHDWGSALGFHWAKRNPERVKGIAFMEFIRP
IPTWDEWPEFARETFQAFRTTDVGRKLIIDQNVFIEGTLPMGVVRPLTEVEMDHYREPFL
NPVDREPLWRFPNELPIAGEPANIVALVEEYMDWLHQSPVPKLLFWGTPGVLIPPAEAA
RLAKSLPNCKAVDIGPGLNLLQEDNPDLIGSEIARWLSTLEISGAPGFSSISAHHHHHHHH
HH
Green: Strep-tag, Red: SNAP-tag, Brown: EGFP, Blue: HaloTag, Purple: His-tag

*Cell Culture and Transfection*

HeLa, U-2 OS (both ATCC), COS-7 (Gift from Dr. R. Sprengel, MPI for Medical Research) or U2OS Nup96-Halo (generously provided by the Ellenberg lab, EMBL) cells were cultured in high-glucose phenol red free DMEM (Life Technologies) medium supplemented with GlutaMAX (Life Technologies), sodium pyruvate (Life Technologies) and 10% FBS (Life Technologies) in a humidified 5% $CO_2$ incubator at 37 °C. Cells were split every 3–4 days or at confluency. These cell lines were regularly tested for mycoplasma contamination. Cells were seeded on glass bottom 35 mm dishes (Mattek or Greiner bio-one), 10 well glass bottom dishes (Greiner bio-one) or 24 mm high precision round coverslips #1.5 (Carl Roth GmbH) one day before imaging. Transient transfection of cells was performed using Lipofectamine™ 2000 reagent (Life Technologies) according to the manufacturer's recommendations: DNA (2.5 µg) was mixed with OptiMEM I (100 µL, Life Technologies) and Lipofectamine™ 2000 (6 µL) was mixed with OptiMEM I (100 µL). The solutions were incubated for 5 min at room temperature, then mixed and incubated for additional 20 min at room temperature. The prepared DNA-Lipofectamine complex was added to a glass bottom 35 mm dish with cells at 50–70% confluency. After 12 h incubation in a humidified 5% $CO_2$ incubator at 37 °C the medium was changed to fresh medium. The cells were incubated for 24–48 h before imaging.

*Stable Cell Line Establishment*

The Flp-In™ T-REx™ System (ThermoFisher Scientific) was used to generate stable cell lines exhibiting tetracycline-inducible expression of the gene of interest (GOI). Briefly, pcDNA5-FRT-TO-GOI and pOG44 were co-transfected into the host cell line U-2 OS FLpIn TREx[11]. Homologous recombination between the FRT sites in pcDNA5-FRT-TO-GOI and on the host cell chromosome, catalysed by the Flp recombinase expressed from pOG44, produced the U-2 OS FlpIn TREx cells expressing stable and inducible the GOI. Selection was performed using 100 µg/mL hygromycin B (ThermoFisher Scientific) and 15 µg/mL blasticidine (ThermoFisher Scientific). Stable cell lines were seeded on glass bottom dishes as described in the previous section, and induced using 100 µg/mL doxycycline (Sigma Aldrich) for 24–48 h previous to imaging.

*Staining*

Cells were stained with 0.2–1 µM PA-SiR (1–2 h, 37 °C) in phenol-red free DMEM medium supplemented with GlutaMAX, sodium pyruvate and 10% FBS (all Life Technologies), washed with the same medium or PBS (once for 3 min, 37 °C) and imaged in the same medium.

*Sample Preparation for Super-Resolution Microscopy*

U-2 OS-CEP41-Halo

U-2 OS-CEP41-Halo cells were seeded on 24 mm glass coverslips and stained with PA-SiR as described above. Methanol fixation was performed as follows: growth medium was removed, cells were incubated for 7 min in –20 °C cold methanol and washed twice with PBS. Fixed-cell samples were mounted in PBS on cavity slides (VWR™) sealed with twinsil® 22 (Picodent) and imaged therein.

F-actin measurements in COS-7 cells

COS-7 cells were seeded on 24 mm glass coverslips and stained with **PA-SiR-Actin** as described above. The cells were fixed as previously described[12]. Briefly, they were fixed and extracted for 1 min using a solution of 0.3% [w/v] glutaraldehyde and 0.25% [v/v] Triton X-100 in CB buffer (CB: 10 mM MES, pH 6.1, 150 mM NaCl, 5 mM EGTA, 5 mM glucose and 5 mM $MgCl_2$), and then post-fixed for 10 min in 2% [w/v] glutaraldehyde in CB. They were treated with freshly prepared 0.1% sodium borohydride for 7 min. Short additional post-staining was performed with 0.5 μM **PA-SiR-Actin** (1 h, 25 °C). Fixed-cell samples were mounted in PBS on cavity slides (VWR™) sealed with twinsil® 22 (Picodent) and imaged therein.

U-2 OS TOMM20-Halo and U-2 OS β-2-adrenergic-receptor-Halo

U-2 OS cells were seeded on 24 mm glass coverslips and stained with **PA-SiR-Halo** as described above. The cells were transiently transfected and the next day the coverslips were mounted into attofluor cell chambers (Life technologies) and the imaging medium was supplemented with HEPES (20 mM). Cells were directly imaged after mounting.

U-2 OS-NUP96-Halo

Genome-edited U-2 OS cells with Halo-tagged NUP96[13] were seeded on 24 mm round coverslips (No. 1.5H; 117640; Marienfeld). Cells were cultured under adherent conditions at 37°C, 5% $CO_2$ and 100% humidity in DMEM (high-glucose, without phenol red) supplemented with 10% [v/v] FBS, 2 mM l-glutamine, nonessential amino acids, and ZellShield. Before sample preparation, the respective dye was added to the medium to a final concentration of 1 μM and incubated for 2 hours. All following incubations were carried out at room temperature and all incubations longer than 1 minute were performed on an orbital shaker in the dark to prevent preactivation of the dye. Cells were prefixed in 2.4% [w/v] formaldehyde (FA) in PBS for 30 seconds, permeabilized in 0.4% [v/v] Triton X-100 in PBS for 3 minutes and fixed in 2.4% [w/v] FA in PBS for 30 minutes. Subsequently, the FA was quenched by incubating the coverslip for 5 minutes in 100 mM $NH_4Cl$ in PBS. After washing 3 times for 5 minutes each in PBS, the coverslips were mounted and imaged in PBS.

*Single-Molecule Assay*

Sample preparation was adapted from two literature procedures[14-15]. Briefly, 18 x 18 mm high precision coverslips (Carl Roth) were sonicated for 10 min in MQ water, 10 min in acetone, 10 min in MeOH, 10 min in KOH (1 M, prepared from 99.98% purity Carl Roth) and rinsed with MQ water after each step. The coverslips were cleaned with piranha solution (1:3, $H_2O_2/H_2SO_4$) twice for 30 min. After extensive rinsing with MQ water they were dried under a $N_2$ stream. A solution of 2% [v/v] *N*-[3-(trimethoxysilyl)propyl]ethylendiamine (Sigma-Aldrich) in dry acetone was prepared and the clean coverslips where immersed in the dark for 1 h. The coverslips were rinsed with acetone, MQ water and then dried with $N_2$. A solution of 1 mg biotin-PEG-SVA (MW 5000, Laysan Bio) and 54 mg mPEG-SVA (MW 5000, Laysan Bio) was prepared in 230 µL sodium bicarbonate buffer (10 mM freshly prepared) and applied to three coverslip pairs. After 3 h in the dark the coverslips were washed with MQ water, blow dried with $N_2$ and stored under $N_2$ at –20 °C. Flow chambers were assembled at need from one glass slide (Carl Roth) and one coated coverslip separated by double sided tape and fixed with epoxy glue.

100 µL of a 0.2 mg/mL solution of streptavidin (Life Technologies) in PBS was applied to the flow chamber and incubated for 10 min. The channel was washed with 400 µL PBS. A solution of SNAP-tag:EGFP:HaloTag (5 µM), fluorophore (2.5 µM), biotin-ligand (5 µM; SNAP-Biotin™ (NEB), HaloTag Biotin (Promega)), in PBS was prepared and incubated for 1 h. 100 µL of a 1:1000–1:500 dilution thereof was applied to the flow chamber and incubated for 10 min. The channel was washed with 400 µL PBS and filled with PBS. They were imaged in TIRF mode using a Leica SR GSD (Supplementary Table S9).

*Widefield Microscopy*

Imaging was performed using a Leica DMi8 microscope (Leica Microsystems) equipped with a Leica DFC9000 GT sCMOS camera; a CoolLED Pe4000 LED light source (635 nm, 635/18; 470 nm, 474/27; 365 nm, 378/52); a HC PL APO 40.0x1.10 water objective and standard GFP (515/40) and Cy5 (720/100) filter sets. Activation of the fluorophores was achieved by irradiation with the 365 nm LED and the DAPI filter set (430/35) at 100% LED output for the indicated durations. The microscope was equipped with a $CO_2$ and temperature controllable incubator (PeCon, 37 °C).

Stability measurement: images were taken in the Cy5 (500 ms, ex: 10%), transmission (100 ms) and the GFP channel (100 ms, ex: 5%) every 30 s. Activation was performed for 1 s once.

Activation experiment: images were taken in the Cy5 (500 ms, ex: 10%), transmission (100 ms) and the GFP channel (100 ms, ex: 5%) consecutively every 9 s. Activation was performed for 50 ms after each acquisition cycle.

*Confocal Microscopy*

Confocal imaging was performed on a Leica DMi8 microscope (Leica Microsystems) equipped with a Leica TCS SP8 X scanhead; a SuperK white light laser, a 355 nm CW laser (Coherent), a HC PL APO 63x1.47 oil objective or a HC PL APO 40.0x1.10 water objective; emission was collected as indicated in Supplementary Table S9. Photoactivation was performed for one frame by using a 355nm laser. The microscope was equipped with a $CO_2$ and temperature controllable incubator (Life Imaging Services, 37 °C).

Signal to Background measurement: cells were focused in the transmission channel and z-stacks were recorded with 0.4 µm step size before and after activation. The summed stacks were analysed as following: the mean of a rectangular ROI within the nucleus was divided by the mean of a rectangular ROI adjacent to the nucleus.

*Super-Resolution Microscopy*

Super-resolution microscopy was performed on a Leica SR GSD (Leica Microsystems) microscope equipped with an Andor iXon3 897 EMCCD camera (Andor) using a central 180 pixel x 180 pixel or 400 pixel x 400 pixel subregion of the camera chip. The system was equipped with the following lasers for excitation and photoactivation: a 642 nm (500 mW; MPBC, Inc.), a 532 nm (1000 mW; MPBC, Inc.), a 488 nm (500 mW; MPBC, Inc.) and a 405 nm (30 mW; Coherent, Inc) diode laser for photoactivation. The standard Leica filter sets for SR GSD systems were used - in brief: Leica set “488” for 405 nm and 488 nm excitation: DBP 405/10 488/10 excitation filter, LP 505 dichroic mirror and 555/100 suppression / emission filter; Leica set “532” for 405 nm and 532 nm excitation: DBP 405/10 532/10 excitation filter, LP 550 dichroic mirror and 600/100 suppression / emission filter; Leica set “642” for 405 nm and 642 nm excitation: DBP 405/10 642/10 excitation filter, LP 650 dichroic mirror and 710/100 suppression / emission filter. Fluorescence was collected through a high-numerical-aperture (NA) oil-immersion objective (Leica HC PL APO 160x1.43). Lateral drift was minimized by the suppressed motion (SuMo) stage of the Leica SR GSD and by keeping the temperature of the environment stable via an incubation box (T = 21 ± 0.1 °C) covering the entire microscope. The microscope was operated by the Leica LAS X software (version 1.9.0.13747). The specific parameters can be found in Supplementary Table S9.

NUP96-Halo samples were imaged on a custom-built epi-fluorescence microscope with homogenous high power illumination[16]. The output of a commercial LightHub laser box (Omicron-Laserage Laserprodukte) with 405 nm, 488 nm, 561 nm and 640 nm laser lines and an additional 640 nm booster laser (Toptica) were focused on a speckle reducer (LSR-3005-17S-VIS; Optotune) and coupled into a multi-mode fiber (M105L02S-A; Thorlabs). The output of this fiber is magnified by an achromatic lens, cleaned up by a quadband filter (390/482/563/640 HC Quad; AHF) and focused into the sample. Fluorescence was collected through a high-numerical-aperture (NA) oil-immersion objective (160x/1.43-NA; Leica), filtered by a 700/100 bandpass filter (AHF) and focused onto an Evolve512D EMCCD camera (Photometrics). The focus was stabilized by a total internally reflected IR laser that was focused onto a quadrant photodiode, which was coupled into a closed-loop with the piezo objective positioner. Typically, we acquire 15,000 to 30,000 frames with 50 ms exposure time and laser power densities of about 13 kW $cm^{-2}$. Data was acquired until no more activated fluorophores were observed. The pulse-length of the 405 nm laser was adjusted during the acquisition to maintain a similar number of localizations per frame. The different components of the microscope are managed by a field-programmable gate array (Mojo; Embedded Micro) which is controlled using a custom-written plugin for µManager[17].

*Software and Image Processing*

Statistical analysis as well as curve fitting was performed using OriginLab[18]. All images except the NUP96-Halo images were processed with ImageJ/Fiji[19-20]. Super-resolution images and TIRF data from the single-molecule assay were processed with the ImageJ plugin ThunderSTORM[21]. Tracking data were analyzed using the TrackMate plugin[22]. Single-molecule assay data were further processed by a costume written MatLab script provided by Dr. Christian Sieben (EPFL) based on the Crocker, Weeks, and Grier Algorithm[23]. Microtubule diameter analysis was performed in ImageJ using the line profile function. The measured lines are indicated in Supplementary Fig. S11a, b (line width = 500 nm). Line profiles were fit with a Gaussian function ($y(x) = y_0 + A /(w \cdot \mathrm{sqrt}(\pi/2)) \cdot e^{-2(x-xc)^\wedge 2/w^\wedge 2}$) and the FWHM was derived (FWHM = sqrt(2 · w ln(2)) · w). Live-cell SMLM data was additionally processes using the HAWK plugin using 3 levels and time grouping, followed by multi-emitter fit in ThunderSTORM allowing for 5 emitters per fitting region[24]. Costume written MatLab code was used to produce the rolling frame video. The movie presented was convoluted with a Gaussian function (sigma = 12 nm).

The reconstruction of super-resolved images of NUP96-Halo was done using the custom-written software SMAP (Superresolution Microscopy Analysis Platform, https://github.com/jries/SMAP). First, localizations were detected using a difference of Gaussians algorithm and a dynamic threshold to exclude random signal fluctuations. Then the localizations were fit by a pixelated Gaussian function. Dim localizations (localization precision > 30 nm) and out-of-focus localizations (fitted size of the Gaussian > 160 nm) were filtered out. Localizations that were found within 75 nm of each other in consecutive frames with maximum 1 frame dark time were grouped into one localization. Images were reconstructed by plotting all localized emitters at the fitted positions as Gaussians with a width proportional to their localization precision.

### Supplementary Figures and Tables

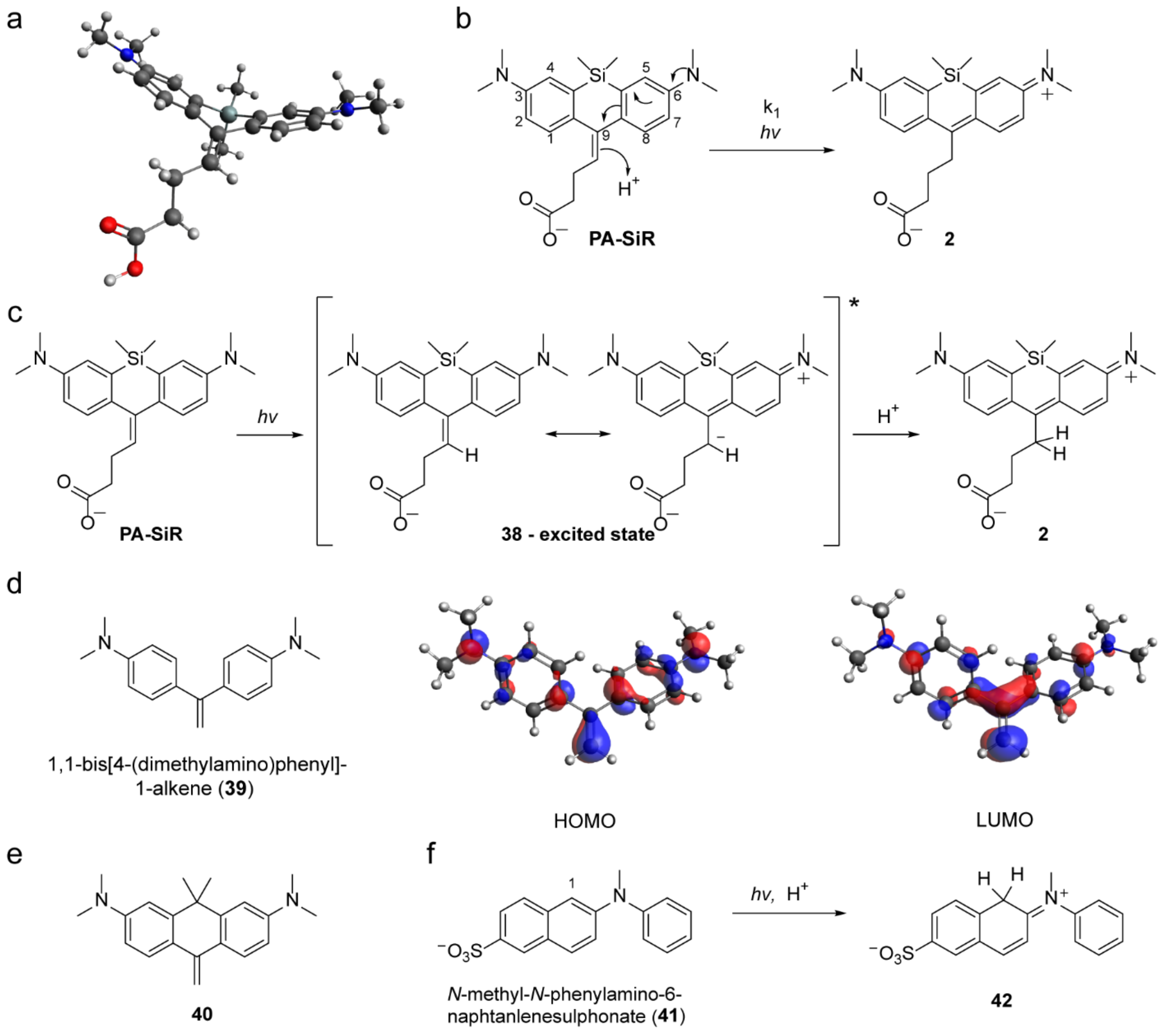


**Supplementary Figure S1.** Photoactivation reaction mechanism (**a**) Optimized structure of **PA-SiR** showing a non-planar conformation of the aromatic system. (**b**) Scheme of the photo-induced isomerization-protonation of **PA-SiR**. (**c**) Light-induced charge transfer leads to excited state **38**, which is protonated and forms **2**. (**d**) Structure along with calculated HOMO and LUMO of model compound 1,1-bis[4-(dimethyl-amino)phenyl]-1-alkene (**39**). Substituted 1,1-diphenylethenes are known to exhibit twisted intramolecular charge transfer states in polar solvents and are therefore comparable to PA-SiRs[25]. The most closely related 1,1-bis[4-(dimethyl-amino)phenyl]-1-alkene shown here is reported to convert to a colored species upon irradiation, but the phenomenon was not further investigated[26]. Its LUMO receives dominant contributions from the extra-cyclic double bond, whereas the HOMO is localized on the periphery of the molecule similarly to PA-SiR **4**. (**e**) Structure of the only reported olefinic rhodamine derivative: carbopyronine **40**[27]. They were hardly isolable and not reported photoactivatable. (**f**) Isolated report on the isomerization and protonation of *N*-methyl-*N*-phenylamino-6-naphtanlenesulphonate (**41**) to protonated **42**[28-29].

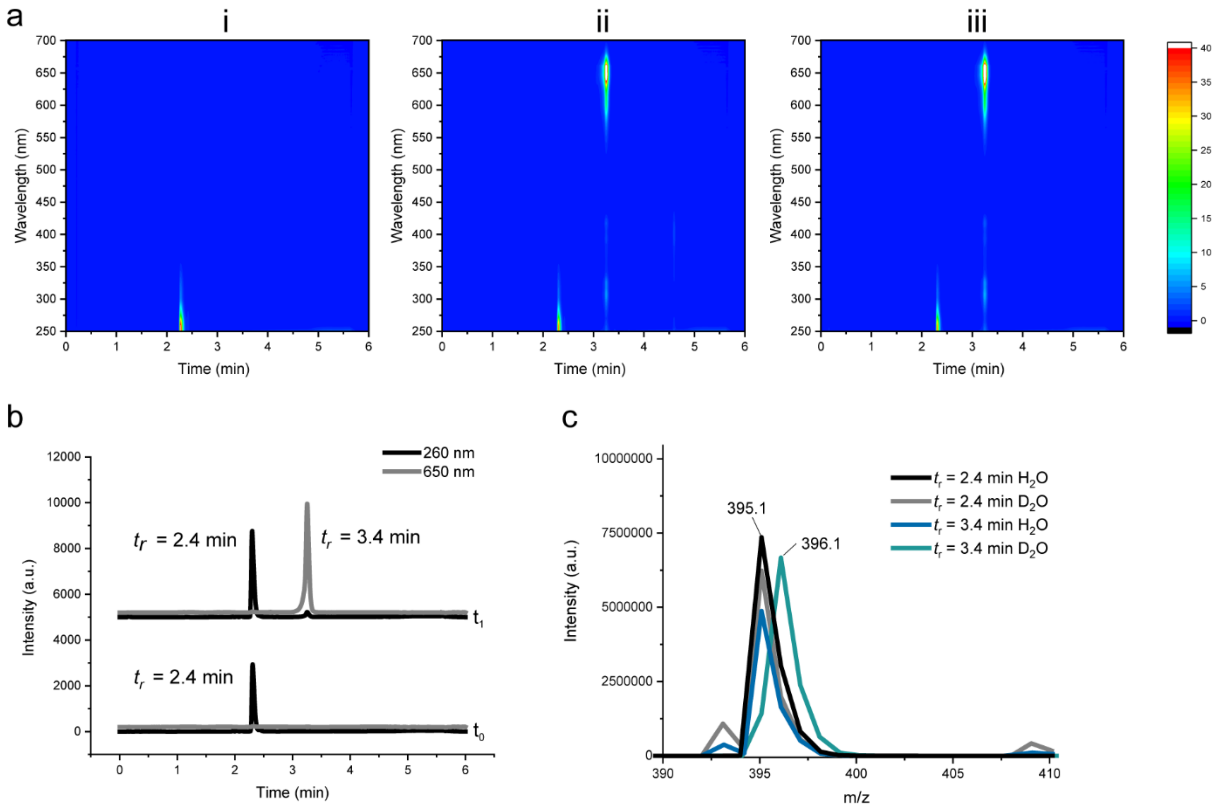


**Supplementary Figure S2**. LC-MS analysis of the photoactivation. (**a**) LC-MS 3D contour plots of aqueous **PA-SiR** solution before (i), directly after (ii) and further 15 min in the dark after activation ensuring complete decay and formation of **3** (iii). It is not possible to observe **3** under the acidic LC-MS conditions. Only **PA-SiR** and SiR **2** are observed (peaks at 2.4 min and 3.4 min). A minor impurity formed during the reaction is visible at 4.8 min (*m/z* = 325.1, 366.1). (**b**) Isotope experiment: LC-MS traces of aqueous ($D_2O$) **PA-SiR** solutions before ($t_0$) and after activation with UV light for 30 s ($t_1$) at 260 nm and 650 nm. Retention times ($t_r$) are given. (**c**) Relevant *m/z* signals of the two peaks in both $H_2O$ and $D_2O$ experiments. The peak at 2.4 min shows the same *m/z* = 395.1 ratio corresponding to **PA-SiR** $[M+H]^+$ in both solvents ($H_2O$ and $D_2O$). The peak at 3.4 min in $D_2O$ has a *m/z* = 396.1 corresponding to deuterated SiR **2** $[M]^+$ confirming protonation/deuteration.

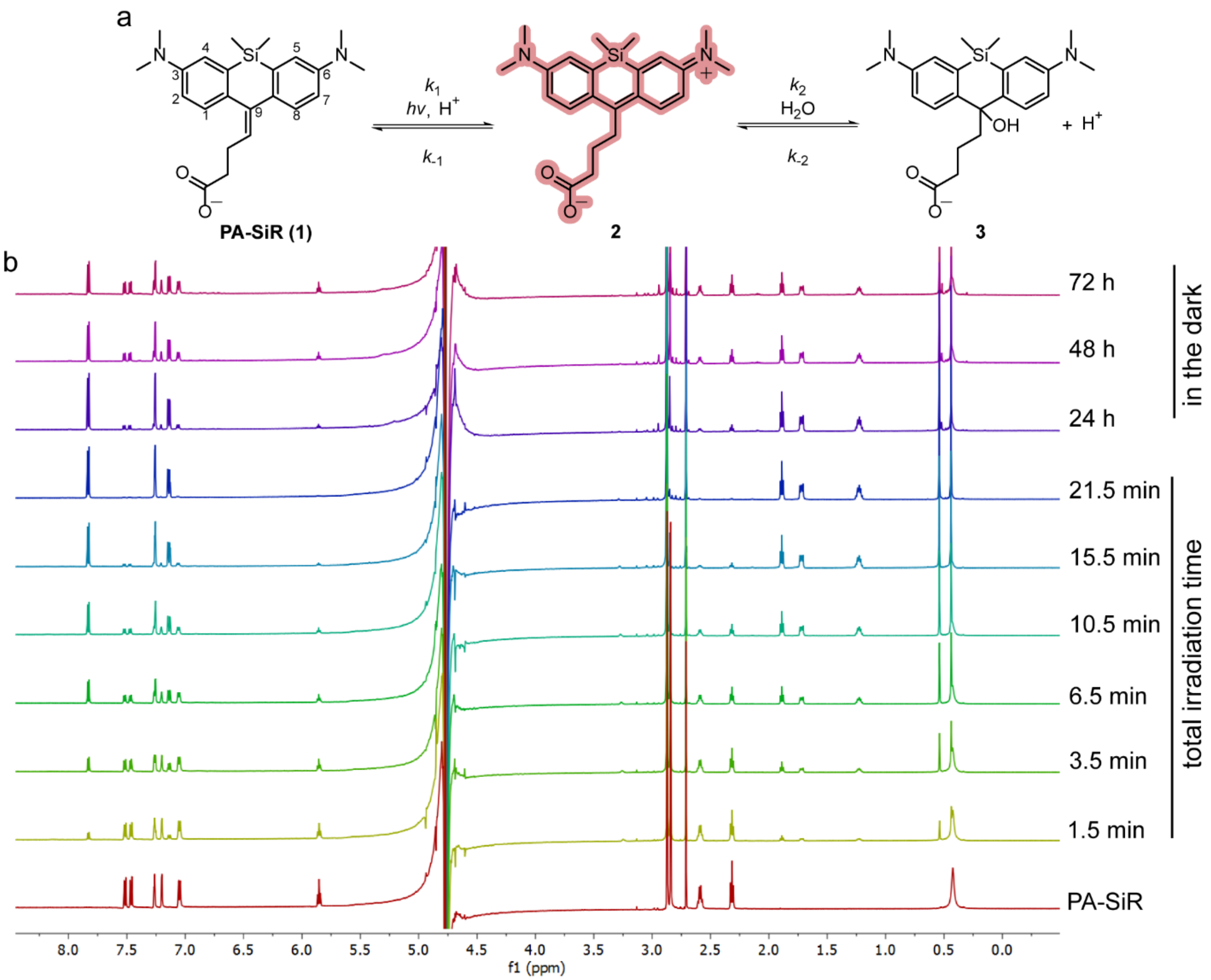


**Supplementary Figure S3.** $^{1}$H nuclear magnetic resonance (NMR) photoactivation experiments (**a**) Equilibrium system of **PA-SiR**, SiR **2** and **3**. (**b**) $^{1}$H NMR spectra of **PA-SiR** (2.0 mM in PBS) before, during and after UV irradiation. Irradiation times are given as total irradiation times. Assignment of the pure spectra of **PA-SiR** and at 21.5 min of **3**: $^{1}$H NMR (**PA-SiR**, 600 MHz, $H_2O + D_2O$): δ 7.52 (d, J = 8.5 Hz, 1H; C$H_{ar}$), 7.46 (d, J = 8.5 Hz, 1H; C$H_{ar}$), 7.26 (d, J = 2.8 Hz, 1H; C$H_{ar}$), 7.20 (d, J = 2.8 Hz, 1H; C$H_{ar}$), 7.06 (d, J = 1.7 Hz, 1H; C$H_{ar}$), 7.04 (d, J = 1.6 Hz, 1H; C$H_{ar}$), 5.85 (t, J = 7.4 Hz, 1H; C*H*), 2.87 (s, 6H; N*Me*), 2.84 (s, 6H; N*Me*), 2.71 (s, 2H; DMSO reference), 2.59 (q, J = 7.5 Hz, 2H; C$H_2$), 2.32 (t, J = 7.5 Hz, 2H; C$H_2$), 0.43 (s, 6H; Si$Me_2$); and $^{1}$H NMR (21.5 min **3**, 600 MHz, $H_2O + D_2O$): δ 7.83 (d, J = 8.8 Hz, 2H; C$H_{ar}$), 7.26 (d, J = 2.8 Hz, 2H; C$H_{ar}$), 7.14 (dd, J = 8.9, 2.8 Hz, 2H; C$H_{ar}$), 2.87 (s, 12H; N$Me_2$), 2.71 (s, 3H; DMSO reference), 1.89 (t, J = 7.4 Hz, 2H; C$H_2$), 1.66 – 1.80 (m, 2H; C$H_2$), 1.12 – 1.29 (m, 2H; C$H_2$), 0.54 (s, 3H; Si*Me*), 0.44 (s, 3H; Si*Me*). The signal around 4.7 ppm corresponds to the residual $H_2O$ signal that was not suppressed. **2** is not visible under the chosen conditions pH= 7–8. This experiments shows that the photoactivation reaction is reversible over a longer time scale, where either $k_{-1}$ or a third process $k_3$ becomes important.

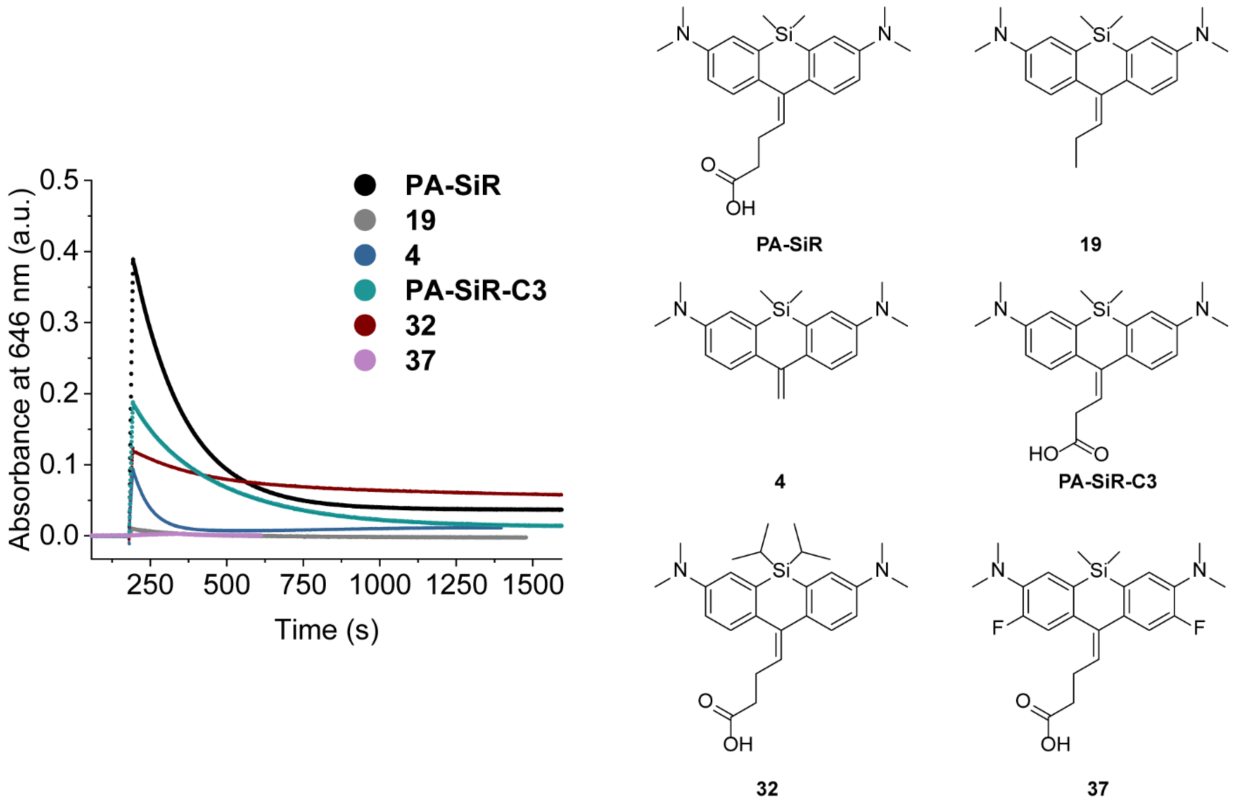


**Supplementary Figure S4**. Time dependent absorbance measurements at 646 nm for different PA-SiR analogues carried out at 10 μM in PBS. Activation was performed for 12 s after 3 min. These measurements let us conclude that PA-SiR analogues are indeed attacked by an external nucleophile, as the removal of the carboxylic acid group of **PA-SiR**, leading to compounds **19** and **4**, did not prevent the nucleophilic attack. Upon structural variation, both faster and slower reaction rates $k_2$ and different equilibrium positions relative to **PA-SiR** were found. For instance, fluorination at the aromatic core lead to PA-SiR derivatives, which only showed a change in absorbance after prolonged UV irradiation. Bulky $^{i}$Pr groups on Si drastically decreased the reaction rates of the nucleophilic attack $k_2$.. Fitted decay parameters are given in the Supplementary Table S6.

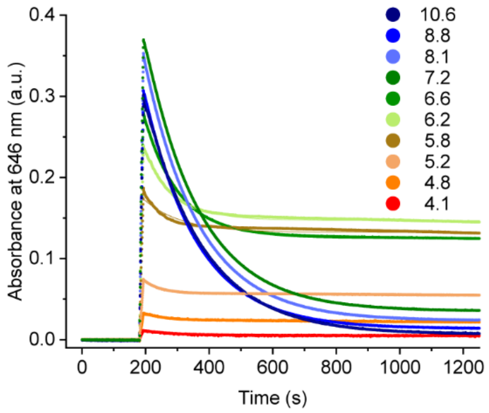


**Supplementary Figure S5.** Influence of different pH values on the equilibrium system. Absorbance measurements at 646 nm over time for **PA-SiR** solutions at 10 μM in PBS with pH values ranging from 4.1–10.6. These measurements were used to extract the normalized $A_{max}$ and $A_{eq}$ (normalized relative to $A_{max}$) values presented in Fig. 1d. All data were fitted with a mono-exponential decay function. The curves obtained at pH values between 6.6 and 4.8 show a bi-exponential decay, indicating the presence of a second component. It is likely that the reaction $k_{-1}$ becomes relevant under these conditions. Therefore, considering exclusively the second equilibrium (represented by $k_2$ and $k_{-2}$) for the fits is only an approximation.

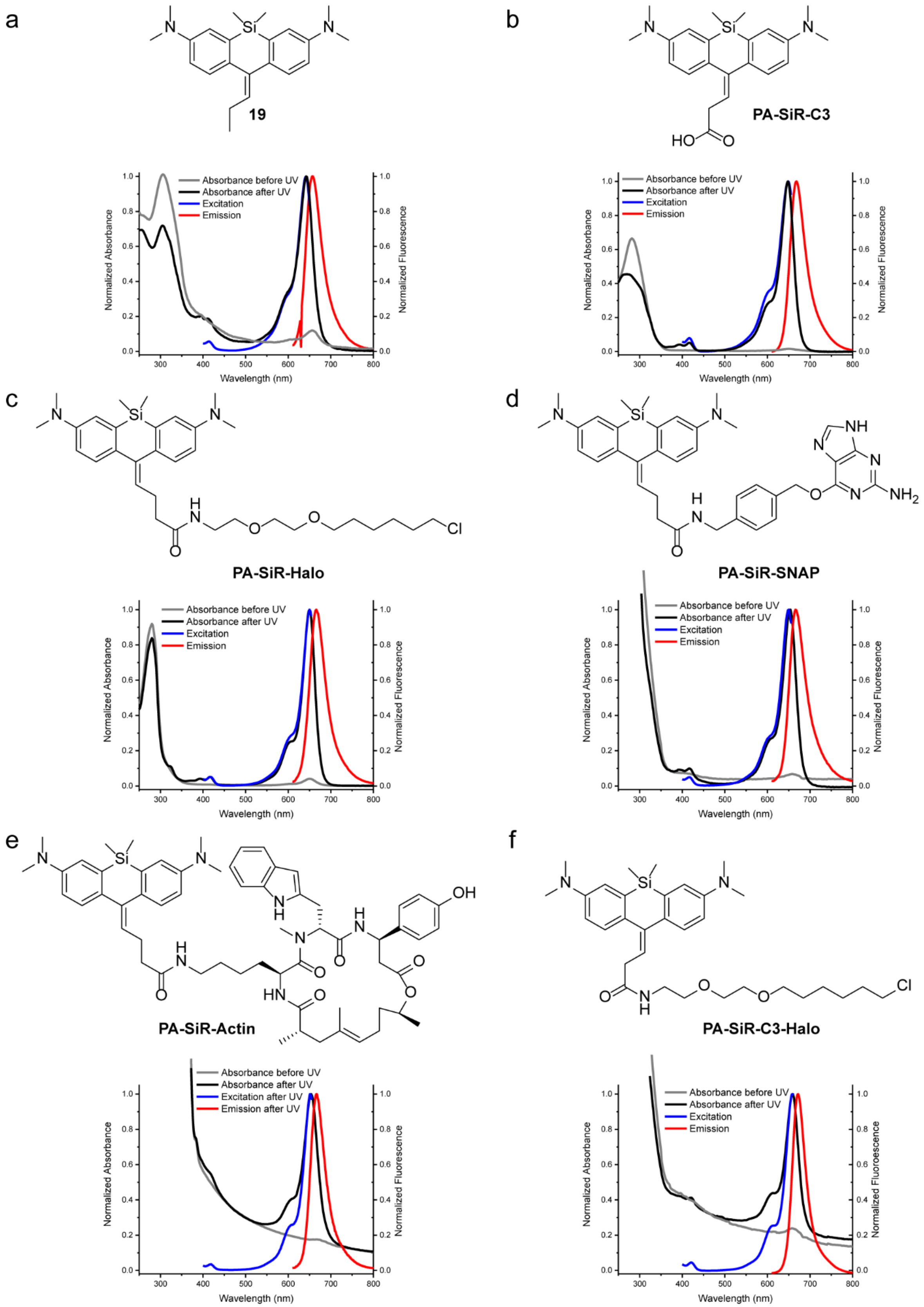

a
19
b
PA-SiR-C3
c
PA-SiR-Halo
d
PA-SiR-SNAP
e
PA-SiR-Actin
f
PA-SiR-C3-Halo
Absorbance before UV
Absorbance after UV
Excitation
Emission
Excitation after UV
Emission after UV
Normalized Absorbance
Normalized Fluorescence
Normalized Fluoroescence
Wavelength (nm)

**Supplementary Figure S6.** Structures and absorption, excitation and emission spectra of PA-SiR derivatives. (**a**) **19**, (**b**) **PA-SiR-C3**, (**c**) **PA-SiR-Halo** conjugated to HaloTag, (**d**) **PA-SiR-SNAP** conjugated to SNAP-tag, (**e**) **PA-SiR-Actin** containing F-actin, (**f**) **PA-SiR-C3-Halo** conjugated to HaloTag. Absorption spectra were measured before and after UV irradiation. All spectra were normalized to their maximum between 600 and 700 nm except the absorbance spectra before UV irradiation, which was normalized to the same factor as the spectra recorded after UV irradiation.

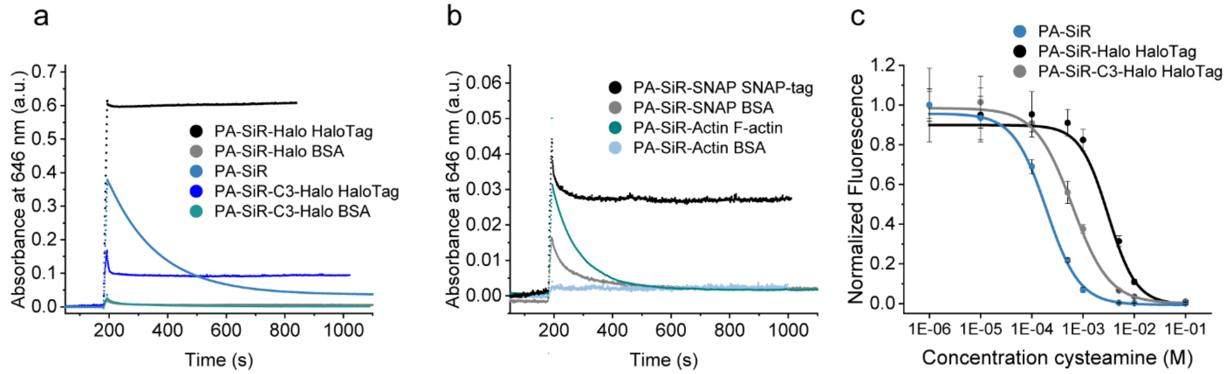


**Supplementary Figure S7.** Absorbance measurements at 646 nm over time for different PA-SiR probes at 10 µM in PBS. Activation was performed for 12 s after 3 min. (**a**) PA-SiR-Halo analogues were measured in the presence of BSA or HaloTag (20 µM). Reaction with HaloTag influences not only the photoactivation but also the kinetics and thermodynamics of the second equilibrium. **PA-SiR-C3-Halo** bearing one methylene group less than **PA-SiR-Halo** showed a fast initial decay after photoactivation followed by a stable signal. In addition, the shortening of the linker was accompanied by a decrease in extinction coefficient (Supplementary Table S1, S4). (**b**) **PA-SiR-SNAP** and **PA-SiR-Actin** were measured with the addition of BSA, SNAP-tag (20 µM) or F-actin, respectively. In comparison to the HaloTag probes, **PA-SiR-SNAP** showed less distinct differences in activation and stability between BSA and SNAP-tag addition conditions. In both cases, nucleophilic attack was observed but the signal of **PA-SiR-SNAP** conjugated to SNAP-tag decayed to a lesser degree, remaining to a higher degree in the fluorescent form. **PA-SiR-Actin** did not activate at all in the presence of BSA. Once bound to F-actin it activated better but still decayed over time. This behavior could be due to the probe unbinding from F-actin and subsequently decaying in solution. The structures of the probes investigated in (**a**) and (**b**) are given in Supplementary Fig. S6. (**c**) Fluorescence signal after addition of cysteamine (0.001–100 mM) to fully activated **PA-SiR**, **PA-SiR-Halo** and **PA-SiR-C3-Halo** conjugated to HaloTag (1 µM dye on 2 µM HaloTag) solutions in equilibrium. $EC_{50}$ (half maximal effective concentration) is determined as: 0.192 ± 0.019 mM for **PA-SiR** (mean ± 95% confidence interval, all $N = 24$ samples), 3.1 ± 0.5 mM for **PA-SiR-Halo** conjugated to HaloTag and 0.62 ± 0.06 mM for **PA-SiR-C3-Halo** conjugated to HaloTag.

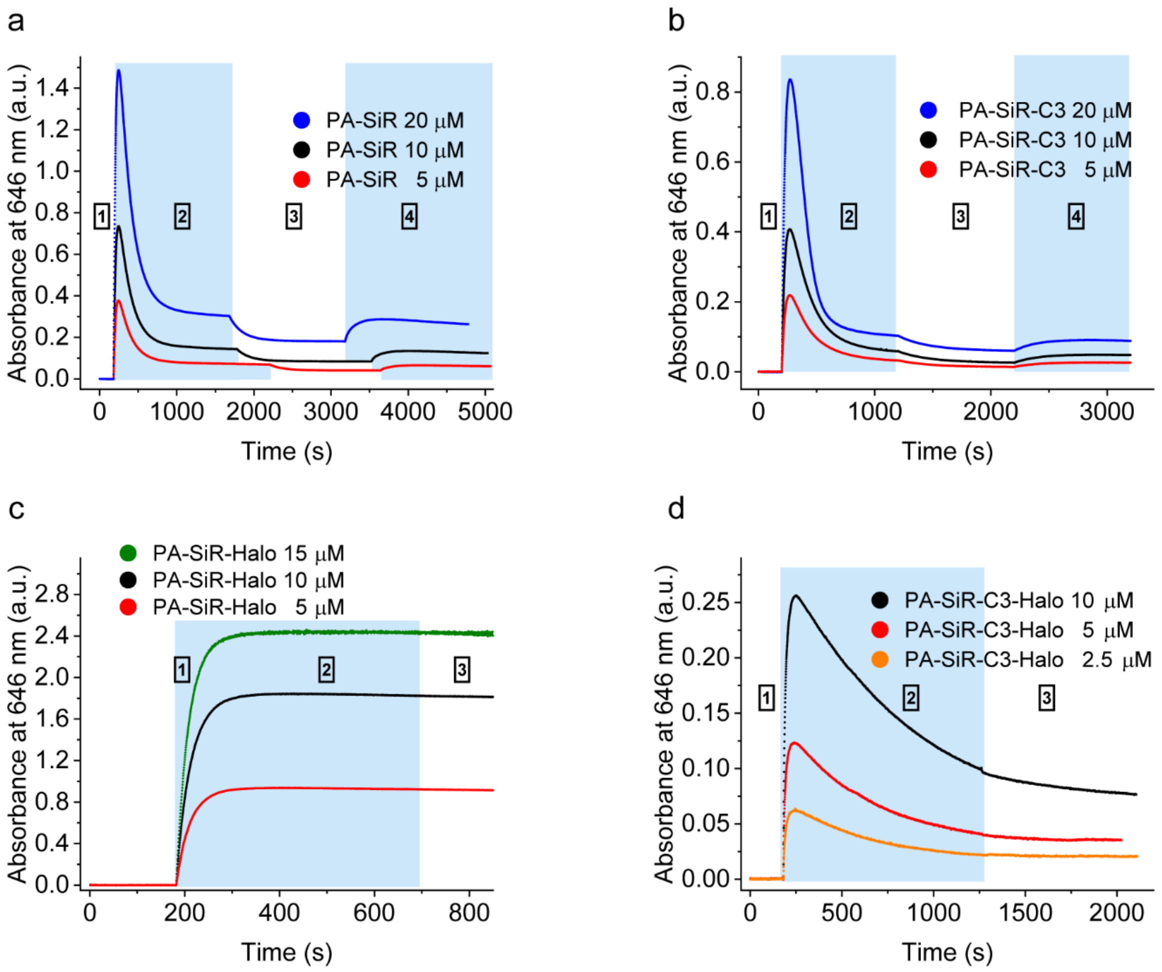


**Supplementary Figure S8**. Saturation experiment of PA-SiRs at different concentrations. (**a**) **PA-SiR**. (**b**) **PA-SiR-C3**. (**c**) **PA-SiR-Halo**. (**d**) **PA-SiR-C3-Halo**. Samples were continuously UV irradiated (light blue color) during the first (increase in absorbance) and second section (decrease in absorbance). UV irradiation was discontinued in the third and restarted in the fourth section. Section one was fitted with a mono-exponential increase to investigate the kinetic parameter $k_1$ of the photoactivation reaction. Section 3 was fitted with a mono-exponential decay and the fitted parameters were used to calculate the theoretical extinction coefficient for the photoproducts (Supplementary Table S1, S7-8).

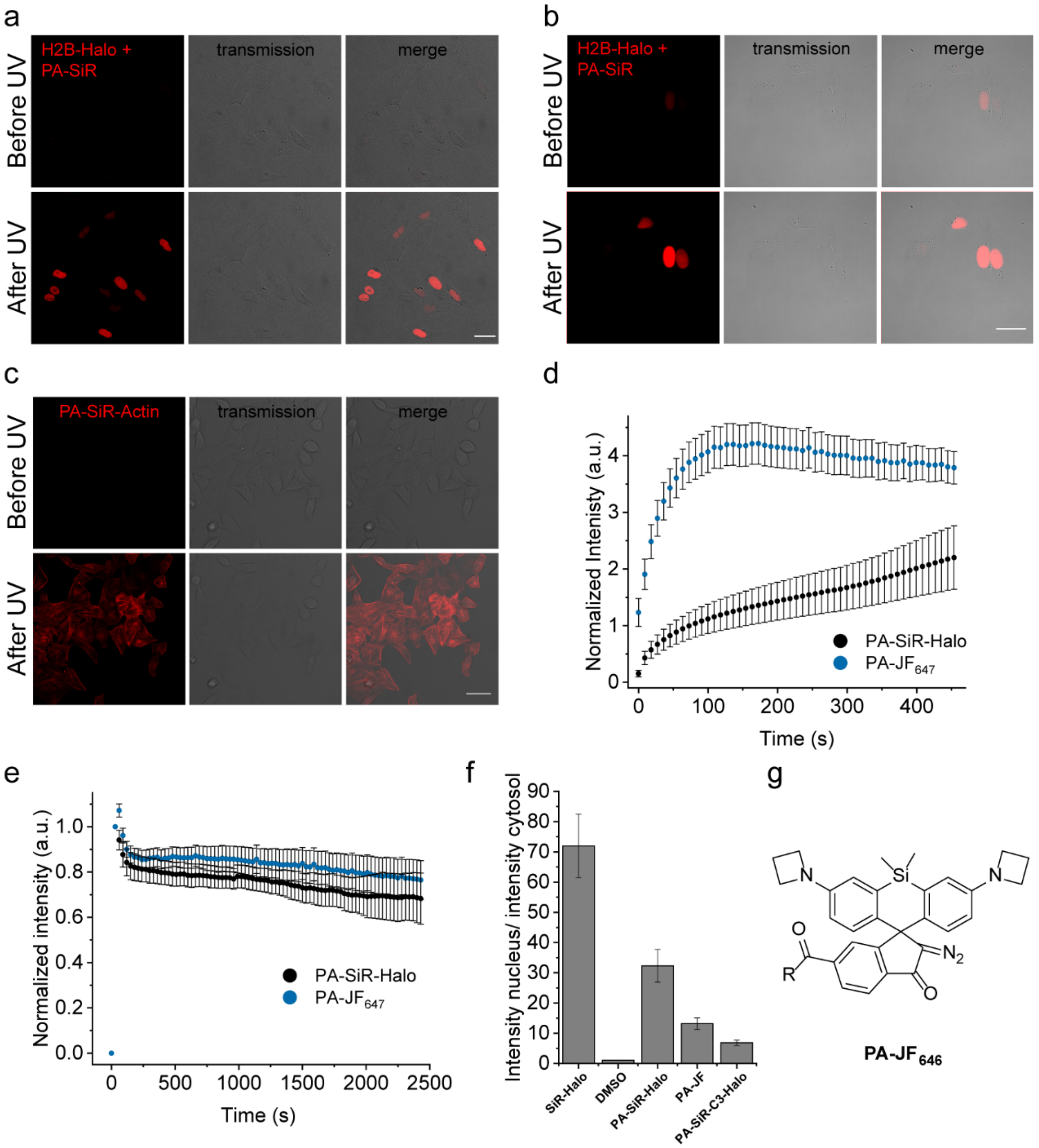


**Supplementary Figure S9.** Confocal and widefield microscopy images (**a-b**) Microscopy images of U-2 OS cells transiently expressing H2B-Halo before and after activation with UV light. Stained with **PA-SiR-Halo** (0.5 µM for 2 h) and activated with the DAPI channel (430/35 nm) on a widefield microscope (**a**) or the 355 nm laser on a confocal set up (**b**). Scale bars, 40 µm. (**c**) Confocal images of HeLa cells stained with **PA-SiR-Actin** (1 µM for 1.5 h) before and after UV activation. Scale bar, 40 µm. (**d**) Activation profiles of **PA-SiR-Halo** and **PA-JF$_{646}$** normalized to the GFP signal in U-2 OS cells transiently expressing H2B-Halo-T2A-EGFP stained with the respective dye (0.5 µM for 2 h). One field of view (FOV) is repeatedly activated with UV light on a widefield set-up (mean ± 95% confidence interval, $N$ = 26 cells for **PA-SiR-Halo**, $N$ = 54 cells for **PA-JF$_{646}$-Halo**). **PA-SiR-Halo** shows a 14-fold turn on whereas **PA-JF$_{646}$**

shows only a 3.5-fold turn on. (**e**) Stability of fluorescence signal over time after activation of **SiR-Halo** and **PA-JF$_{646}$-Halo** localized to H2B-Halo-T2A-EGFP (mean ± 95% confidence interval, *N* = 30 cells for **PA-SiR-Halo** and 70 cells for **PA-JF$_{646}$**). (**f**) Signal-over-background measurements for different fluorophores (250 nM for 1 h) in U-2 OS cells expressing H2B-Halo-T2A-EGFP (mean ± 95% confidence interval): **SiR-Halo** (72 ± 11, *N* = 135 cells), DMSO (1.04 ± 0.07, *N* = 126 cells), **PA-SiR-Halo** (32 ± 5, *N* = 119 cells), **PA-JF$_{646}$-Halo** (13.2 ± 1.9, *N* = 121 cells), **PA-SiR-C3-Halo** (6.8 ± 0.9, *N* = 86 cells), all after activation. (**g**) Chemical structure of **PA-JF$_{646}$**.

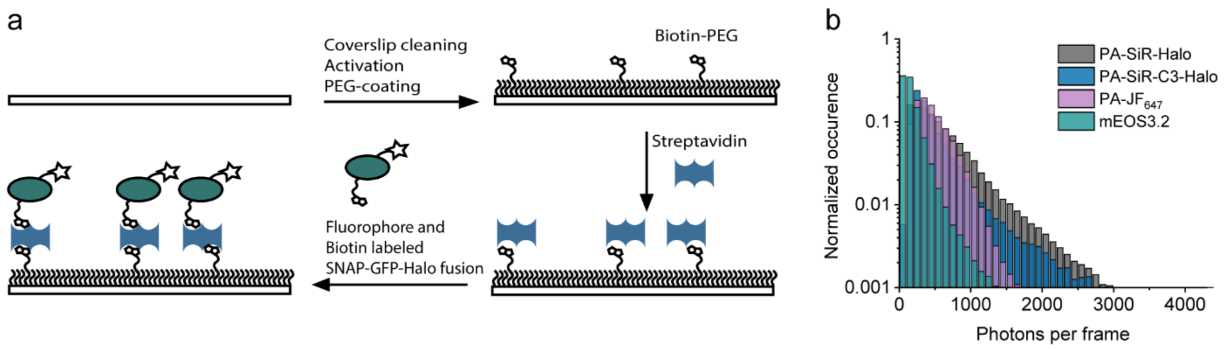


**Supplementary Figure S10**. Single-molecule assay. (**a**) Schematic view of the *in vitro* TIRF fluorophore assay: A coverslip is cleaned, the surface activated with an amino-silane and then PEG coated. The PEG-brush contains sparse PEG-biotin moieties for fluorophore immobilization. First, PEG-biotin is saturated with streptavidin followed by SNAP:EGFP:Halo or mEOS3.2:Halo labeled with either biotin-BG and fluorophore-chloroalkane or biotin-chloroalkane. (**b**) Photons per particle per frame for **PA-SiR-Halo**, **PA-SiR-C3-Halo** and **PA-JF$_{646}$-Halo** at 642 nm (1.2 kW cm$^{-2}$, live-cell tracking regime) and mEOS3.2 at 532 nm (0.9 kW cm$^{-2}$), respectively. **PA-SiR-Halo** (668, 518) showed a 30% higher photon output as **PA-SiR-C3-Halo** (497, 348) and **PA-JF$_{646}$-Halo** (474, 414) (mean, median). **mEOS3.2** (187, 132), on the other hand, showed much lower photon numbers then the small-molecule fluorophores, as expected from literature.[30]

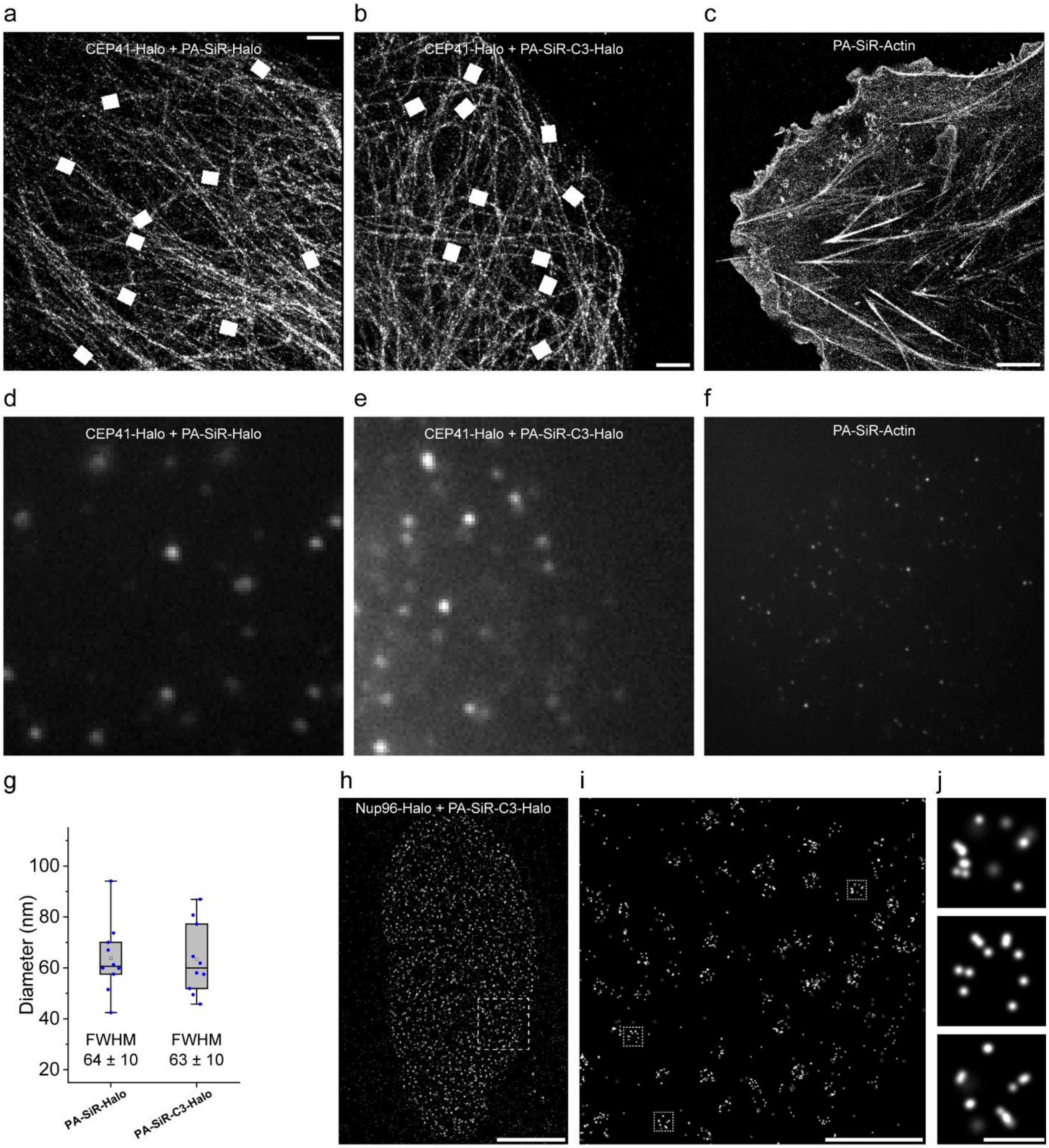


**Supplementary Figure S11.** Fixed-cell SMLM and quantification. (**a**) Super-resolved image from Figure 3a with the indicated areas for microtubule diameter analysis. Scale bar, 1 µm. (**b**) Super-resolved image of a fixed U-2 OS cell, stably expressing CEP41-Halo stained with **PA-SiR-C3-Halo** (1 µM for 2 h), reconstructed from 6004 frames (100 ms exposure time, 4.2 kW cm$^{-2}$ at 642 nm excitation). Even though the performance of **PA-SiR-C3-Halo** was not optimal in live-cells (Supplementary Fig. S9f), it was possible to obtain good SMLM images after fixation and washing. The microtubule diameter was found to be FWHM$_{\textbf{PA-SiR-C3-Halo}}$ = 63 ± 10 nm ($N$ = 10 tubules). The areas for microtubule diameter analysis are indicated. Scale bar, 1 µm. (**c**) Super-resolved image of fixed COS-7 cells stained with **PA-SiR-Actin** (0.5 µM for 2 h). Scale bar, 5 µm. (**d-f**) Single raw camera frames from the recordings to give the super-resolved images above. (**g**) Box-plots of the distribution for the FWHM obtained by fitting the intensity profiles measured along the lines indicated in **a** and **b** (line profiles): box = 25%–75% percentile, whiskers 5%–95% percentile, black line = median, box = mean, individual data points in blue. (**h**) Super-resolved overview image of endogenously tagged Nup96-Halo in U-2 OS cells stained with **PA-SiR-C3-Halo** (1 µM for 2 h). Scale bar, 5 µm. (**i**) Super-resolved image from the boxed region in **h**. Scale bar, 1 µm. (**j**) Single nuclear pores from boxed regions in **i** following the same order (top-bottom). Scale bar, 100 nm.

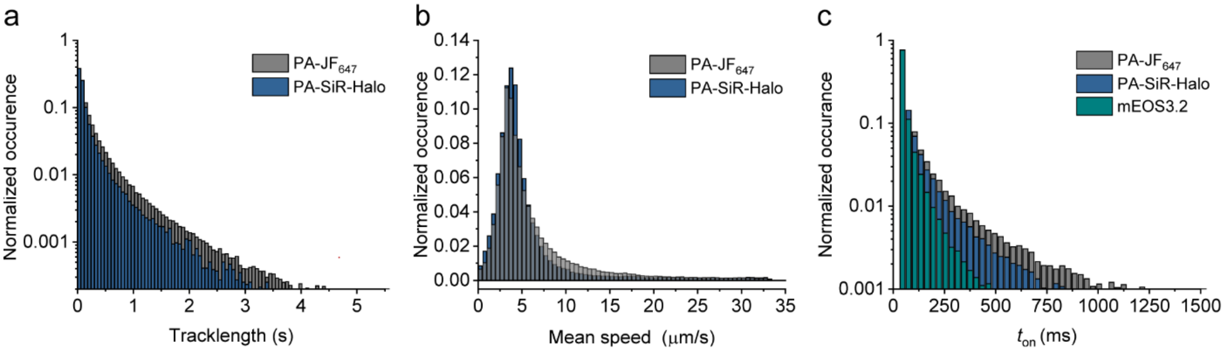


**Supplementary Figure S12.** Quantification of live-cell tracking experiments in U-2 OS cells. (**a**) Histograms of tracklength found for **PA-SiR-Halo** (208, statistical mean; no deviation given as not normally distributed; $N$ = 51'408 tracks from 2 FOV) and **PA-JF$_{646}$-Halo** (338, $N$ = 233'871 tracks from 2 FOV) labeling β-2-adrenergic-receptor-Halo under identical imaging conditions (30 ms, 0.3 kW cm$^{-2}$ at 642 nm). **PA-SiR-Halo** displayed similar tracklengths as **PA-JF$_{646}$-Halo** and is therefore well suited for live-cell tracking experiments. (**b**) Determined mean speed for **PA-SiR-Halo** (5.55 ± 0.05 μm/s, median = 4.1 μm/s) and **PA-JF$_{646}$-Halo** (6.36 ± 0.02 μm/s, median = 4.4 μm/s) (mean ± 95% confidence interval, median). (**c**) On-times found in the single-molecule assay for **PA-SiR-Halo** and **PA-JF$_{646}$-Halo** at 642 nm (1.2 kW cm$^{-2}$, live-cell tracking regime) and mEOS3.2 at 532 nm (0.9 kW cm$^{-2}$), respectively, showing that the photostability for mEOS3.2 is considerably lower than for the two small-molecule fluorophores.

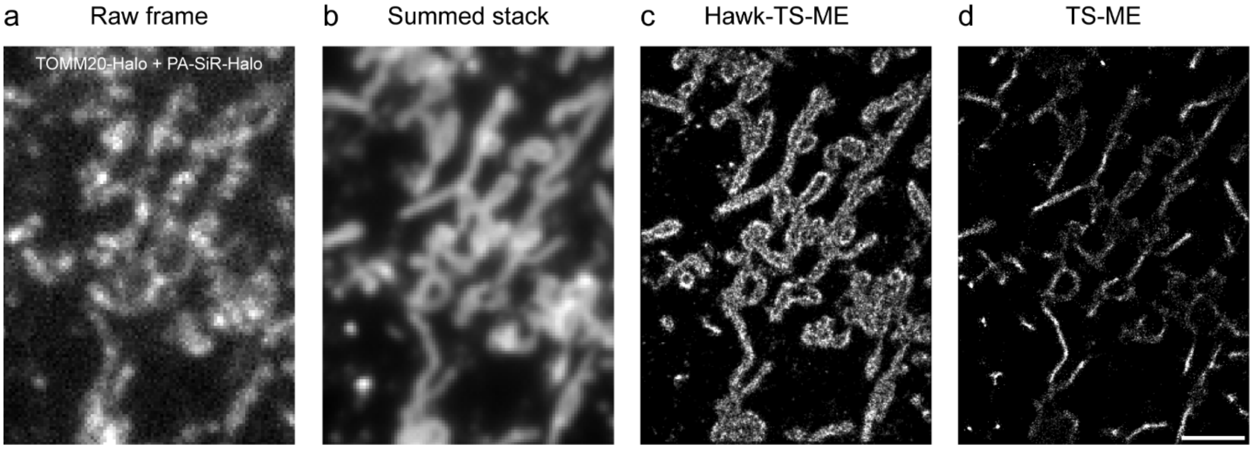


**Supplementary Figure S13.** Live-cell SMLM experiment. (**a**) Single raw camera frame. (**b**) Sum projection over the first 10 s as depicted in Figure 4b. (**c**) Super-resolved image acquired at high-emitter-density within 10 s (50 ms, 0.3 kW $cm^{-2}$ at 642 nm) from Figure 4a. Processed with the HAWK ImageJ plugin[24] and ThunderSTORM multi-emitter fit[21]. (**d**) Super-resolved image obtained if the data is only processed by ThunderSTORM multi-emitter fit[21], resulting in artificial narrowing and collapsing of structures. Scale bar, 2 μm.

**Supplementary Table S1.** Spectral properties of different PA-SiR analogues. Results are given as means ± 95% confidence interval. Except for the quantum yield of activation, which is a propagated standard error of the mean. *N* = number of samples measured.

| | PA-SiR before photoactivation | | | Photoproduct | | | | |
|---|---|---|---|---|---|---|---|---|
| # | $\lambda_{abs,max}$ [nm] | $\varepsilon_{max}$ [$M^{-1}cm^{-1}$] | $\varphi_{act}$ % | $\lambda_{ex,\ max}$/ $\lambda_{em,\ max}$ [nm] | $\varepsilon_{max,\ 646\ nm}$ [$M^{-1}cm^{-1}$] | $\varphi$ % | $\tau_{PBS\ or\ BSA}$ [s] | $\tau_{on\ POI}$ [s] |
| **PA-SiR** | 276 | 17'000±3'000 *N* = 10 | 13.3±2.3 | 646/664 | 90'000±18'000 *N* = 3 | 19.0±2.4 *N* = 4 | 169±9 *N* = 3 | N/A |
| **SiR-COOH** | N/A | N/A | N/A | 645/661*,** | 100'000** | 39** | N/A | N/A |
| **19** | 314 | 16'000±5'000 *N* = 9 | N/D | 642/656 | N/D | 13±10 *N* = 3 | 200±70 *N* = 3 | N/A |
| **4** | N/D | N/D | N/D | N/D | N/D | 11±8 *N* = 3 | 55±21 *N* = 3 | N/A |
| **PA-SiR-C3** | 281 | 16'000±3'000 *N* = 9 | 29±13 | 648/668 | 39'000±7'000 *N* = 3 | 15.6±1.5 *N* = 3 | 271±15 *N* = 3 | N/A |
| **32** | N/D | N/D | N/D | N/d | N/D | 16.1±0.7 *N* = 3 | 460±60 *N* = 3 | N/A |
| **PA-SiR-Halo** | 333 | 33'000±5'000 *N* = 9 | 0.86±0.07 | 650/666 | 180'000±30'000 *N* = 3 | 29.2±1.2 *N* = 3 | 40±60 *N* = 3 | 6±3 *N* = 3 |
| **PA-SiR-C3-Halo** | 329 | 20'000±3'000 *N* = 9 | 3.9±0.3 | 658/672 | 15'000±3'000 *N* = 3 | 29±5 *N* = 3 | 100±18 *N* = 3 | 7±4 *N* = 3 |
| **PA-SiR-SNAP** | 294 | 19'000±6'000 *N* = 10 | N/D | 650/666 | N/D | 11.1±2.0 *N* = 3 | 84±19 *N* = 3 | 30±50 *N* = 3 |
| **PA-SiR-Actin** | 298 | 21'000±7'000 *N* = 9 | N/D | 651/666 | N/D | 3±6 *N* = 3 | no act | 92±3 *N* = 3 |

$\lambda_{abs,max}$ absorption maximum, $\varepsilon_{max}$ extinction coefficient, $\varphi_{act}$ quantum yield of activation, $\lambda_{ex,\ max}$/ $\lambda_{em,\ max}$ excitation and emission maximum, $\varphi$ quantum yield, $\tau$ and $\tau_{on\ POI}$ mean lifetime, N/D not determined, N/A not applicable, * $\lambda_{abs,max}$/ $\lambda_{em,\ max}$, ** from [8].

**Supplementary Table S2**. Fit parameters from Gaussian fit in Fig. 1d. $\mathbf{y(x) = y_0 + A /(w \cdot sqrt(\pi/2)) \cdot e^{-2(x-xc)^2/w^2}}$

| Name | $y_0$ [a.u.] | $se_{y0}$ [a.u.] | xc | $se_{xc}$ | w | $se_w$ | A | $se_A$ | $X^2$ | $R^2$ |
|---|---|---|---|---|---|---|---|---|---|---|
| $A_{eq}$ | 0.000 | 0.095 | **5.339** | 0.115 | 1.972 | 0.283 | 2.564 | 0.560 | 16.507 | 0.984 |

$y_0$ offset and its standard error $se_{y0}$; **xc** center and its standard error $se_{xc}$; **w** width and its standard error $se_w$; **A** area and its standard error $se_A$; $X^2$ the reduced chi-squared and $R^2$ the squared correlation coefficient are given.

**Supplementary Table S3**. Fit parameters from sigmoidal fit in Fig. 1d. $\mathbf{y(x) = a/(1 + e^{-k(x-xc)})}$

| Name | a [a.u.] | $se_a$ [a.u.] | xc | $se_{xc}$ | k | $se_k$ | $X^2$ | $R^2$ |
|---|---|---|---|---|---|---|---|---|
| $A_{max}$ | 0.900 | 0.032 | **5.731** | 0.084 | 2.465 | 0.435 | 0.004 | 0.971 |

**a** amplitude and its standard error $se_a$; **xc** center and its standard error $se_{xc}$; **k** coefficient and its standard error $se_k$; $X^2$ the reduced chi-squared and $R^2$ the squared correlation coefficient are given.

**Supplementary Table S4.** Averaged fit parameters from mono-exponential fits for PA-SiR probes in Fig. 2b and Supplementary Fig. S7a, b. Results are given as means and standard deviations from three measurements. $\mathbf{y(t) = y_0 + A \cdot e^{(-(t-x)/\tau)}}$

| | Fit Parameters | | | | | | Calculated | | | | | | |
|---|---|---|---|---|---|---|---|---|---|---|---|---|---|
| **Name** | $\tau$ [s] | $sd_\tau$ [s] | A [a.u.] | $sd_A$ [a.u.] | $y_0$ [a.u.] | $sd_{y0}$ [a.u.] | $A_{eq}$ [a.u.] | $A_{x=0}$ [a.u.] | $K_2$ | $k_{app}$ [$s^{-1}$] | $k_{-2}$ [$s^{-1}$] | $k_2$ [$s^{-1}$] | $K_{-2}$ |
| **PA-SiR-Halo BSA** | 41.98 | 23.46 | 0.012 | 0.001 | 0.006 | 0.002 | 0.006 | 0.017 | 1.99 | 2.38E-02 | 7.97E-03 | 1.59E-02 | 0.50 |
| **PA-SiR-Halo Halo** | 5.74 | 1.19 | 0.013 | 0.004 | 0.593 | 0.013 | 0.593 | 0.606 | 0.02 | 1.74E-01 | 1.70E-01 | 3.87E-03 | 43.99 |
| **PA-SiR-C3-Halo BSA** | 105.02 | 7.34 | 0.018 | 4.4E-04 | -0.000* | 0.001 | 0.000 | 0.018 | N/D | 9.52E-03 | N/D | N/D | N/D |
| **PA-SiR-C3-Halo Halo** | 6.51 | 1.55 | 0.062 | 0.002 | 0.095 | 0.002 | 0.095 | 0.157 | 0.66 | 1.54E-01 | 9.27E-02 | 6.09E-02 | 1.52 |
| **PA-SiR-SNAP BSA** | 84.10 | 7.49 | 0.011 | 0.001 | 0.002 | 0.001 | 0.002 | 0.014 | 4.68 | 1.19E-02 | 2.09E-03 | 9.80E-03 | 0.21 |
| **PA-SiR-SNAP SNAP** | 32.66 | 18.20 | 0.010 | 0.003 | 0.026 | 0.007 | 0.026 | 0.036 | 0.37 | 3.06E-02 | 2.23E-02 | 8.35E-03 | 2.67 |
| **PA-SiR-Actin Actin** | 92.45 | 1.35 | 0.032 | 0.005 | 0.002 | 0.002 | 0.002 | 0.035 | 14.50 | 1.08E-02 | 6.98E-04 | 1.01E-02 | 0.07 |

$\tau$ decay constant and its standard deviation $\mathbf{sd_\tau}$; **A** amplitude and its standard deviation $\mathbf{sd_A}$; $y_0$ offset and its standard deviation $\mathbf{sd_{y0}}$, along with the derived parameters: $\mathbf{A_{eq} = y_0}$; $\mathbf{A_{x=0} = A + A_{eq}}$; $\mathbf{K_2 = (A_{x=0} - A_{eq}) \cdot A_{eq}^{-1}}$; $\mathbf{k_{app} = \tau^{-1}}$; $\mathbf{k_{-2} = k_{app} \cdot (K_2 + 1)^{-1}}$; $\mathbf{k_2 = k_{-2} - k_{app}}$; $\mathbf{K_{-1} = k_2 \cdot k_{-2}^{-1}}$**.** *negative equilibrium values are within the experimental error of the instrument and were set to 0 for further calculations. N/D not determined.

**Supplementary Table S5.** Fit parameters from the dose response fit in Fig. 2c and Supplementary Fig. S7c. **y(x) = A1 + (A2-A1)/(1 + 10^((LOGx0-x)*p))**

| Name | A1 | $se_{A1}$ | A2 | $se_{A2}$ | Logx0 | $se_{Logx0}$ | p | $se_p$ | EC50 | $se_{EC50}$ | $X^2$ | $R^2$ |
|---|---|---|---|---|---|---|---|---|---|---|---|---|
| **PA-SiR** | -0.0064 | 0.0083 | 0.9568 | 0.0103 | -3.7167 | 0.0205 | -1.3449 | 0.0647 | **1.92E-04** | 9.04E-06 | 0.001 | 0.997 |
| **PA-SiR-Halo** | -0.0043 | 0.0207 | 0.8989 | 0.0119 | -2.5127 | 0.0326 | -1.5812 | 0.1251 | **3.07E-03** | 2.30E-04 | 0.001 | 0.991 |
| **PA-SiR-C3-Halo** | 0.0013 | 0.0108 | 0.9842 | 0.0097 | -3.2101 | 0.0186 | -1.2019 | 0.0699 | **6.16E-04** | 2.64E-05 | 0.001 | 0.997 |

**A1** and **A2** asymptotes and their standard errors $\mathbf{sd_{A1}}$ and $\mathbf{sd_{A2}}$; **LOGx0** center and its standard error $\mathbf{se_{LOGx0}}$; **p** hill slope and its standard error $\mathbf{se_p}$; **EC50** derived parameter concentration at half response and its standard error $\mathbf{se_{EC50}}$; $X^2$the reduced chi-squared and $R^2$ the squared correlation coefficient are given.

**Supplementary Table S6.** Averaged fit parameters from mono-exponential fits in Supplementary Fig. S4 and S5. Results are given as means and standard deviations from three measurements. $\mathbf{y(t) = y_0 + A \cdot e^{(-(t-x)/\tau)}}$

| | Fit Parameters | | | | | | Calculated | | | | | | |
|---|---|---|---|---|---|---|---|---|---|---|---|---|---|
| **Name/pH** | $\tau$ [s] | $sd_\tau$ [s] | A [a.u.] | $sd_A$ [a.u.] | $y_0$ [a.u.] | $sd_{y0}$ [a.u.] | $A_{eq}$ [a.u.] | $A_{x=0}$ [a.u.] | $K_2$ | $k_{app}$ [$s^{-1}$] | $k_{-2}$ [$s^{-1}$] | $k_2$ [$s^{-1}$] | $K_{-2}$ |
| **PA-SiR PBS** | 169.50 | 3.49 | 0.347 | 0.008 | 0.035 | 0.002 | 0.035 | 0.382 | 9.87 | 5.90E-03 | 5.43E-04 | 5.36E-03 | 0.10 |
| **19** | 202.87 | 28.89 | 0.009 | 0.001 | -0.002* | 0.001 | 0.000 | 0.009 | N/D | 4.93E-03 | N/D | N/D | N/D |
| **4** | 54.92 | 8.49 | 0.086 | 0.002 | 0.008 | 0.004 | 0.008 | 0.094 | 10.25 | 1.82E-02 | 1.62E-03 | 1.66E-02 | 0.10 |
| **PA-SiR-C3** | 271.19 | 6.02 | 0.172 | 0.003 | 0.013 | 0.001 | 0.013 | 0.185 | 13.21 | 3.69E-03 | 2.59E-04 | 3.43E-03 | 0.08 |
| **32** | 460.09 | 22.57 | 0.057 | 0.011 | 0.050 | 0.007 | 0.050 | 0.106 | 1.14 | 2.17E-03 | 1.02E-03 | 1.16E-03 | 0.88 |
| **10.6** | 180.19 | 8.60 | 0.283 | 0.014 | 0.004 | 0.005 | 0.004 | 0.287 | 74.93 | 5.55E-03 | 7.31E-05 | 5.48E-03 | 0.01 |
| **8.8** | 171.83 | 5.29 | 0.303 | 0.021 | 0.010 | 0.006 | 0.010 | 0.313 | 30.75 | 5.82E-03 | 1.83E-04 | 5.64E-03 | 0.03 |
| **8.1** | 171.83 | 4.12 | 0.310 | 0.021 | 0.019 | 0.006 | 0.019 | 0.329 | 16.63 | 5.82E-03 | 3.30E-04 | 5.49E-03 | 0.06 |
| **7.2** | 169.55 | 2.85 | 0.322 | 0.045 | 0.033 | 0.004 | 0.033 | 0.355 | 9.65 | 5.90E-03 | 5.54E-04 | 5.34E-03 | 0.10 |
| **6.6** | 114.99 | 0.65 | 0.150 | 0.003 | 0.121 | 0.003 | 0.121 | 0.271 | 1.23 | 8.70E-03 | 3.90E-03 | 4.80E-03 | 0.81 |
| **6.2** | 96.94 | 2.78 | 0.094 | 0.007 | 0.146 | 0.001 | 0.146 | 0.240 | 0.64 | 1.03E-02 | 6.27E-03 | 4.04E-03 | 1.55 |
| **5.8** | 93.18 | 22.73 | 0.043 | 0.004 | 0.130 | 0.004 | 0.130 | 0.173 | 0.33 | 1.07E-02 | 8.08E-03 | 2.65E-03 | 3.05 |
| **5.2** | 97.69 | 26.62 | 0.016 | 0.002 | 0.057 | 0.002 | 0.057 | 0.072 | 0.28 | 1.02E-02 | 8.02E-03 | 2.22E-03 | 3.62 |
| **4.8** | 115.09 | 65.13 | 0.009 | 0.001 | 0.020 | 0.002 | 0.020 | 0.028 | 0.45 | 8.69E-03 | 6.01E-03 | 2.68E-03 | 2.24 |
| **4.1** | 114.50 | 7.21 | 0.006 | 4.2E-04 | 0.005 | 2.3E-04 | 0.005 | 0.010 | 1.16 | 8.73E-03 | 4.05E-03 | 4.68E-03 | 0.87 |

$\boldsymbol{\tau}$ decay constant and its standard deviation $\mathbf{sd_\tau}$; **A** amplitude and its standard deviation $\mathbf{sd_A}$; $\boldsymbol{y_0}$ offset and its standard deviation $\mathbf{sd_{y0}}$, along with the derived parameters: $\mathbf{A_{eq} = y_0}$; $\mathbf{A_{x=0} = A + A_{eq}}$; $\boldsymbol{K_2 = (A_{x=0} - A_{eq}) \cdot A_{eq}^{-1}}$; $\boldsymbol{k_{app} = \tau^{-1}}$; $\boldsymbol{k_{-2} = k_{app} \cdot (K_2 + 1)^{-1}}$; $\boldsymbol{k_2 = k_{-2} - k_{app}}$; $\boldsymbol{K_{-1} = k_2 \cdot k_{-2}^{-1}}$. *negative equilibrium values are within the experimental error of the instrument and were set to 0 for further calculations. N/D not determined.

**Supplementary Table S7.** Fit parameters from mono-exponential fits of the third section for **PA-SiR**, **PA-SiR-C3** (**C3**) and **PA-SiR-C3-Halo** (**C3Halo**) from the saturation experiment in Supplementary Figure S8. Equilibrium values for **PA-SiR-Halo** (**Halo**) were taken from the fit parameters from the first section and are displayed here for completeness. The concentration is given in µM. Results are given for one measurement with the standard error of the fit. $\mathbf{y(t) = y_0 + A \cdot e^{(-(t\text{-}x)/\tau)}}$

| | **Fit Parameters** | | | | | | **Calculated** | | | | | | | | |
|---|---|---|---|---|---|---|---|---|---|---|---|---|---|---|---|
| **Name** | $\tau$ [s] | se$_\tau$ [s] | A [a.u.] | se$_A$ [a.u.] | $y_0$ [a.u.] | se$_{y0}$ [a.u.] | $A_{eq}$ [a.u.] | $A_{x=0}$ [a.u.] | $K_2$ | $k_{app}$ [s$^{-1}$] | $k_{-2}$ [s$^{-1}$] | $k_2$ [s$^{-1}$] | $K_{-2}$ | $K$ | $A_{sat}$ [a.u.] |
| **PA-SiR/5** | 171.26 | 0.16 | 0.028 | 1.5E-05 | 0.040 | 3.9E-06 | 0.040 | 0.069 | 0.69 | 5.84E-03 | 3.45E-03 | 2.39E-03 | 1.44 | 9.866 | **0.440** |
| **PA-SiR/10** | 173.14 | 0.16 | 0.057 | 3.3E-05 | 0.084 | 6.8E-06 | 0.084 | 0.141 | 0.68 | 5.78E-03 | 3.43E-03 | 2.35E-03 | 1.46 | 9.866 | **0.911** |
| **PA-SiR/20** | 172.73 | 0.12 | 0.125 | 4.9E-05 | 0.182 | 1.1E-05 | 0.182 | 0.307 | 0.69 | 5.79E-03 | 3.42E-03 | 2.36E-03 | 1.45 | 9.866 | **1.974** |
| **C3/5** | 268.05 | 0.36 | 0.018 | 9.0E-06 | 0.014 | 6.4E-06 | 0.014 | 0.032 | 1.34 | 3.73E-03 | 1.59E-03 | 2.14E-03 | 0.74 | 13.212 | **0.192** |
| **C3/10** | 274.48 | 0.26 | 0.031 | 1.1E-05 | 0.026 | 8.1E-06 | 0.026 | 0.057 | 1.23 | 3.64E-03 | 1.64E-03 | 2.01E-03 | 0.82 | 13.212 | **0.363** |
| **C3/20** | 269.74 | 0.27 | 0.041 | 1.5E-05 | 0.059 | 1.1E-05 | 0.059 | 0.101 | 0.70 | 3.71E-03 | 2.18E-03 | 1.52E-03 | 1.43 | 13.212 | **0.843** |
| **C3Halo/2.5** | 388.55 | 0.79 | 0.044 | 4.0E-05 | 0.020 | 1.9E-05 | 0.020 | 0.064 | 2.25 | 2.57E-03 | 7.93E-04 | 1.78E-03 | 0.45 | 0.657 | **0.032** |
| **C3Halo/5** | 441.00 | 0.69 | 0.091 | 5.6E-05 | 0.032 | 3.5E-05 | 0.032 | 0.123 | 2.82 | 2.27E-03 | 5.93E-04 | 1.67E-03 | 0.35 | 0.657 | **0.053** |
| **C3Halo/10** | 594.43 | 1.30 | 0.196 | 1.4E-04 | 0.065 | 1.3E-04 | 0.065 | 0.261 | 3.02 | 1.68E-03 | 4.19E-04 | 1.26E-03 | 0.33 | 0.657 | **0.108** |
| **Halo/5** | N/A | N/A | N/A | N/A | 0.929 | 1.7E-04 | 0.929 | N/A | N/A | N/A | N/A | N/A | N/A | 0.020 | **0.947** |
| **Halo/10** | N/A | N/A | N/A | N/A | 1.832 | 2.5E-04 | 1.832 | N/A | N/A | N/A | N/A | N/A | N/A | 0.020 | **1.869** |
| **Halo/15** | N/A | N/A | N/A | N/A | 2.432 | 2.5E-04 | 2.432 | N/A | N/A | N/A | N/A | N/A | N/A | 0.020 | **2.480** |

$\boldsymbol{\tau}$ decay constant and its standard deviation **sd**$_\tau$; **A** amplitude and its standard deviation **sd**$_A$; $\boldsymbol{y_0}$ offset and its standard deviation **sd**$_{y0}$, along with the derived parameters: $\mathbf{A_{eq} = y_0}$; $\mathbf{A_{x=0} = A + A_{eq}}$; $\boldsymbol{K_2 = (A_{x=0} - A_{eq}) \cdot A_{eq}^{-1}}$; $\boldsymbol{k_{app} = \tau^{-1}}$; $\boldsymbol{k_{-2} = k_{app} \cdot (K_2 + 1)^{-1}}$; $\boldsymbol{k_2 = k_{-2} - k_{app}}$; $\boldsymbol{K_{-1} = k_2 \cdot k_{-2}^{-1}}$; $\boldsymbol{K}$ = equilibrium constant from the Supplementary Table S4 or S6 for the respective compound; $\mathbf{A_{sat}} = \boldsymbol{K \cdot A_{eq} + A_{eq}}$. These values where used to calculate the extinction coefficients at 646 nm. N/A not applicable.

**Supplementary Table S8.** Averaged fit parameters from the exponential fits for the first sections of the saturation experiments. Results are given as means and standard deviations from three measurements $\mathbf{y(t) = y_0 + A \cdot e^{(-(t-x)/\tau)}}$

| Name | $\tau$ [s] | $sd_\tau$ [s] |
|---|---|---|
| **PA-SiR** | 19.40 | 0.36 |
| **PA-SiR-Halo** | 28.24 | 1.99 |
| **PA-SiR-C3** | 20.99 | 0.80 |
| **PA-SiR-C3-Halo** | 10.32 | 0.72 |

$\boldsymbol{\tau}$ decay constant and its standard deviation $\mathbf{sd_\tau}$.

**Supplementary Table S9** Settings for the different microscopy experiments.

| Image | Label | Ligand | Microscope | Excitation [nm] | Exposure time (Widefield)/ Pixel dwell time (Confocal) | Activation [nm] | Pinhole | Objective | Size | Fixed-live | Emission [nm] | Comment |
|---|---|---|---|---|---|---|---|---|---|---|---|---|
| Fig. 2d, e | H2B-Halo | PA-SiR-Halo | Confocal | 631 | 0.6 µs | 355 | 1 | 40x/1.10 water | 1024x1024 | Live | 777-800 | Max projection |
| Fig. 2f | CEP41-Halo | PA-SiR-Halo | Confocal | 631 | 0.25 µs | 355 | 1 | 40x/1.10 water | 2488x2488 | Live | 751-779 | |
| Fig. 2g | LA-Halo | PA-SiR-Halo | Confocal | 631 | 0.475 µs | 355 | 1 | 40x/1.10 water | 1288x1288 | Live | 751-779 | |
| Fig. 2h | TOMM20-Halo | PA-SiR-Halo | Confocal | 631 | 1.2 µs | 355 | 1 | 40x/1.10 water | 410x714 | Live | 751-779 | |
| Fig. 3a | CEP41-Halo | PA-SiR-Halo | GSD TIRF | 642 642/10 | 100 ms | 405 405/10 | - | 160x/1.43 oil | - | Fixed-MeOH | LP 649 BP 710/100 | 14’083 frames |
| Fig. 3b | β-2-adrenergic-receptor-Halo | PA-SiR-Halo | GSD TIRF | 642 642/10 | 30 ms | 405 405/10 | - | 160x/1.43 oil | - | Live | LP 649 BP 710/100 | 10’046/ 4’000 frames |
| Fig. 3c-e | NUP96-Halo | PA-SiR-Halo | Widefield (custom) | 640 HC Quad | 50 ms | 405 | - | 160x/1.43 oil | - | Fixed-FA | 700/100 | |
| Fig. 4 | TOMM20-Halo | PA-SiR-Halo | GSD TIRF | 642 642/10 | 50 ms | 405 405/10 | - | 160x/1.43 oil | - | Live | LP 649 BP 710/100 | 1’499 frames |
| S9a = 2d,e | H2B-Halo | PA-SiR-Halo | Confocal | 631 | 0.6 µs | 355 | 1 | 40x/1.10 water | 1024x1024 | Live | 777-800 | |
| S9b | H2B-Halo | PA-SiR-Halo | Widefield | 635 | 500 ms | 365 | - | 40x/1.10 water | - | Live | 720/100 | |
| S9c | none | PA-SiR-Actin | Confocal | 631 | 0.225 µs | 355 | 1 | 63x/1.40 oil | 2688x2688 | Live | 751-779 | |
| S9d | H2B-Halo | Respective dye | Widefield | 635 | 500 ms | 365 | - | 40x/1.10 water | - | Live | 720/100 | 50 ms act |
| S9e | H2B-Halo | Respective dye | Widefield | 635 | 500 ms | 365 | - | 40x/1.10 water | - | Live | 720/100 | Stability |
| S9f | H2B-Halo | Respective dye | Confocal | 631 | 0.6 µs | 355 | 1 | 40x/1.10 water | 1024x1024 | Live | 777-800 | Max projection |
| S10b | Halo:EGFP:SNAP | Respective dye | TIRF | 642 642/10 | 30 ms | 405 405/10 | - | 160x/1.43 oil | 400x400 | - | LP 649 BP 710/100 | 10’000-20’000 frames |
| S10b | mEOS3.2:Halo | - | TIRF | 532 532/10 | 30 ms | 405 405/10 | - | 160x/1.43 oil | 400x400 | - | LP 541 BP 600/100 | 10’000-20’000 frames |
| S11a = Fig. 3a | CEP41-Halo | PA-SiR-Halo | GSD TIRF | 642 642/10 | 100 ms | 405 405/10 | - | 160x/1.43 oil | - | Fixed-MeOH | LP 649 BP 710/100 | 14’083 frames |

| | | | | | | | | | | | | |
|---|---|---|---|---|---|---|---|---|---|---|---|---|
| S11b | CEP41-Halo | PA-SiR-C3-Halo | TIRF | 642<br>642/10 | 100 ms | 405<br>405/10 | - | 160x/1.43 oil | - | Fixed-MeOH | LP 649<br>BP 710/100 | 6’004 frames |
| S11c | - | PA-SiR-Actin | GSD<br>TIRF | 642<br>642/10 | 130 ms | 405<br>405/10 | - | 160x/1.43 oil | - | Fixed-Glutaraldehyde | LP 649<br>BP 710/100 | 30’000 frames |
| S11e-g | NUP96-Halo | PA-SiR-C3-Halo | Widefield (custom) | 640<br>HC Quad | 50 ms | 405 | - | 160x/1.43 oil | - | Fixed-FA | 700/100 | |
| S12a,b | β-2-adrenergic-receptor-Halo | Respective dye | GSD<br>TIRF | 642<br>642/10 | 30 ms | 405<br>405/10 | - | 160x/1.43 oil | - | Live | LP 649<br>BP 710/100 | |
| S12c | Halo:EGFP:SNAP | Respective dye | TIRF | 642<br>642/10 | 30 ms | 405<br>405/10 | - | 160x/1.43 oil | 400x400 | - | LP 649<br>BP 710/100 | 10’000-20’000 frames |
| S12c | mEOS3.2:Halo | - | TIRF | 532<br>532/10 | 30 ms | 405<br>405/10 | - | 160x/1.43 oil | 400x400 | - | LP 541<br>BP 600/100 | 10’000-20’000 frames |
| S13a-c | TOMM20-Halo | PA-SiR-Halo | GSD<br>TIRF | 642<br>642/10 | 50 ms | 405<br>405/10 | - | 160x/1.43 oil | - | Live | LP 649<br>BP 710/100 | 1’499 frames |

### Movies

**Supplementary Movie S1.** SMLM rolling frame movie of mitochondria dynamics. U-2 OS cells expressing TOMM20-Halo labeled with PA-SiR-Halo (0.5 μM, 1 h). Each frame is reconstructed from 200 frames (10 s). Scale bar 1 μm.
